# DNMT inhibitors increase methylation at subset of CpGs in colon, bladder, lymphoma, breast, and ovarian, cancer genome

**DOI:** 10.1101/395467

**Authors:** Anil K Giri, Tero Aittokallio

## Abstract

**Background:** DNA methyltransferase inhibitors (DNMTi) decitabine and azacytidine are approved therapies for acute myeloid leukemia and myelodysplastic syndrome. Identification of CpGs violating demethylaion due to DNMTi treatment may help to understand their resistance mechanisms.

**Materials and Methods:** To identify such CpGs, we analysed publicly available 450K methylation data of multiple cancer type cell lines.

*Results:* We identified 637 CpGs corresponding to genes enriched for p53 and olfactory receptor pathways with a transient increase in methylation (median Δβ = 0.12) after decitabine treatment in HCT116 cells. Azacytidine treatment also increased methylation of identified CpGs in 9 colon, 9 ovarian, 3 breast, and 1 lymphoma cancer cell lines.

**Conclusion:** DNMTi treatment increases methylation of subset of CpGs in cancer genome.

## Introduction

DNA methyltransferase inhibitors (DNMTi) are widely used as chemical tools for hypomethylating the genome in order to understand the role of DNA methylations in X-chromosome inactivation, DNA imprinting and transcriptional regulation of several disease-related genes [1-4]. Further, DNMTi agents, decitabine along with its analog azacytidine, have been approved by United States Food and Drug Administration (US FDA), and they currently remain as the sole treatment option for specific sub-groups of acute myeloid leukemia (AML) and myelodysplastic syndrome (MDS) patients [5-6]. Since DNA methylation-induced silencing of tumor suppressor genes, such as P53, at promoter region is a primary event in many cancers and these methylations can be reversed by DNMTi as therapy, both of these drugs are also being tested as a treatment option for breast, lung, colon and other cancers. Decitabine treatment causes global hypomethylation of the genome by intercalating itself in the DNA during replication and halting the DNA methylation transferases (DNMTs) actions [5-6]. Hypomethylation of the genome leads to re-expression of several genes, including multiple tumor suppressor and inhibition of oncogenes, thereby contributing to apoptosis of cancer cells through multiple ways such as DNA damage response pathway, p53 signaling pathways, cytotoxicity, etc [6,7].

However, there are sporadic reports where treatment with DNMTi has led to an increased expression level of DNA methylating enzymes hence DNA methylation in specific cells [8-11]. For example, Kastl *et al*. reported an increase in the mRNA level of DNMT1, DNMT3a and DNMT3b genes in docetaxel-resistant MCF7 cells as compared to drug sensitive cells when treated with decitabine [8]. Surprisingly, a recent study showed that decitabine treatment can cause an increase in 5-hydroxymethylcytosine, an oxidation product of methylated cytosine, in DNA of human leukemic cells [9]. Further, an analog of decitabine, azacytidine treatment, was reported to induce DNA methylation in transgenes of Chinese hamster cell in the process of silencing foreign genes in the human genomes [10]. This piece of evidence hints that treatment with azacytidine can induce DNA methylation at certain locations in the genome that may have non-human origins such as retrotransposons and other genes with viral origin [10,11]. Available piece of literature also suggests that DNMTi treatment causes hypomethylation nearly at 99% of methylated locations in the genome [12], suggesting that there should also be loci where DNMTi treatment can increase the methylation level or has no effect on methylation, instead of the regular role of hypomethylation. However, we are currently lacking the information of the genomic location, function, origin, and fate of those CpGs in the cancer genome that can resist the DNA demethylation.

In the present work, we aim to systematically investigate the extent, location and role of CpGs with increased methylation in response to DNMTi treatment. Identification of such loci and their related genomic features will not only help to understand the reasons behind the failure of the DNMTi treatment in demethylating cancer-related genes but it may also reveal novel molecular mechanism behind efficacy, side effects, and resistance towards DNMTi treatment in various cancer types. We selected HCT116 cell line as our primary disease model to discover these CpGs as it shows the silencing of various tumor suppressor genes due to hypermethylation as seen in the case of colon cancer tissue [13]. Further, HCT116 cell line has been frequently utilized to study DNA methylation and its role in regulating gene expression in colon cancer [14]. To investigate how general these findings are, we tested the increase in methylation after DNMTi treatment identified in HCT116 cells also in other lymphoma, colon, ovarian, and breast cancer cells. Further, we explored the relationship between methylation status of the identified loci and expression status of genes in colon adenocarcinoma cancer using patient tumor data from The Cancer Genome Atlas (TCGA) project. Our work lays foundation for the search of rare events of hypermethylation due to DNMTi treatment contrary to their classic role of DNA hypomethylation in the cancer genome.

## Methodology

### Processing of methylation data

To identify CpGs with increased methylation after decitabine treatment we analyzed the DNA methylation (Illumina 450K platform, GSE51810) and gene expression data (Illumina HumanHT-12_V4_0_R1 platform, GSE51810) from the study by Yang *et al*. [15] for HCT116 colon cell lines treated with decitabine (0.3 mM) for 72 hours. Cells were maintained in McCoy’s 5A medium, supplemented with 10% fetal bovine serum along with 1% penicillin/streptomycin after drug treatment, and followed through 5, 14, 24, 42, and 68 days. The increase in DNA methylation in HCT116 cells were validated using methylation data from the study by Han *et al* [16] (Illumina 450K, GSE41525), where HCT116 and T24 (bladder cancer) cell lines were treated with 0.3 µM and 1 µM of decitabine, respectively for 24 hours and Illumina 450K assay was performed for both untreated and decitabine treated cells. We also tested the increase in DNA methylation of identified CpGs using DMSO (as mock) and decitabine-treated MCF7 cells in data generated by Leadem *et al* (Illumina 450K platform, GSE97483) [17]. These cells were cultured in Minimum Essential Medium (MEM) with 10% fetal bovine serum and treated with 0.06 µM of decitabine for 72 hours.

We also extended our findings discovered in case of decitabine in another DNMTi inhibitor, azacytidine, by analyzing DNA methylation data (Illumina 450K, GSE45707) for untreated and azacytidine-treated (5mM for 72 hours) lymphoma cancer U937 cell line. We further analysed additional methylation data for 26 breast cancer cell lines (MDA231,SKBR3, HCC38, ZR7530, HCC1937, CAMA1, MDA415, HCC1500, BT474, EFM192A, MDA175, MDA468, MDA361, HCC1954, BT20, ZR751, HCC1569, EFM19, T47D, MDA453, MCF7, HCC1187, HCC1419, EFM192A, MDA436, SUM149, and SUM159), 12 colorectal cancer cell lines (SW48, HCT116, HT29, RKO, SW480, Colo320, Colo205, SW620, SNUC-1,CACO-2, SK-CO1, and Colo201), and 13 ovarian cancer cell lines (TykNu, CAOV3, OAW28, OV2008, ES2, EF27, Kuramochi, OVKATE, Hey, A2780, ES2, OVCAR3, OVCAR5, and SKOV3) measured after mock treatment and 0.5 µM azacytidine treatment for 72 hours (Illumina 450K platform, GSE57342). The cells have been cultured and maintained under recommended conditions for each cell line [18].

To investigate the alteration in methylation status of identified probes in cancerous tissue and their role in gene expression regulation, TCGA level 3 HumanMethylation 450K data and normalized RNA-seq gene expression profiles for colon adenocarcinoma (COAD) samples were downloaded using the FireBrowse tool (http://gdac.broadinstitute.org/).

Methylation status at a CpG site was measured as beta value (β) which is the ratio of the methylated probe intensity and the overall intensity (sum of methylated and unmethylated probe intensities designed for a particular CpG in 450K beadchip). It ranges from 0 to 1, indicating no methylation (β=0) to complete methylation of the CpGs (β=1). We performed appropriate quality control of the published data before their downstream analysis. We removed all the CpGs with missing values and a tendency of cross-hybridization as specified in the supplementary file of Chen *et al* [19]. To remove any possible bias due to design differences in the type of probes (the type I and type II probes) present in the Illumina 450K platform, we performed BMIQ normalization [20] to the DNA methylation data for TCGA samples before correlation and differential methylation analysis. All other data processing was done using local inbuilt commands in R as described previously [21].

### Identification of probes with increased methylation in HCT116 cell line

We calculated the difference in methylation level of CpGs before and after treatment with decitabine in HCT116 cell line at day 5 in data from Yang *et al* (GSE51810). CpG that showed an increase in β-value of greater than or equal to 0.10 (Δβ ≥0.10) between untreated control and decitabine treated HCT116 cells after 5 days were identified as CpGs with increased methylations.

### Gene expression data analysis for HCT116 cell line and colon adenocarcinoma tumors from TCGA

Expression analysis was carried out using the inbuilt commands in R. The data were log2-transformed and normalized using Robust Spline Normalization (RSN) using the lumi package in R [22]. Pearson correlation between methylation and gene expression across different time point in HCT116 cell line was assessed using the cor.test function of R.

Gene expression data from TCGA samples were normalized using voom function in the limma package [23], and these data were Z-transformed before the differential and correlation analyses. Wilcoxon non-parametric test was used to identify differentially expressed genes in TCGA adenocarcinoma samples (FDR<0.05). Only those adenocarcinoma samples that had both DNA methylation and gene expression information were used for the correlation analyses.

### Gene annotation and pathway enrichment analysis

Identified CpGs were annotated for their location in the genome based on annotation file provided by Illumina (ftp://ussd-ftp.illumina.com/downloads/ProductFiles/HumanMethylation450/HumanMethylation450_15017482_v1-2.csv). Gene ontology and pathway enrichment analysis of genes corresponding to CpGs with increased methylation were done using GeneCodis [24]. Statistical enrichment was assessed by FDR corrected p-values from hypergeometric test for separate ontology terms and pathways. GENEMANIA [25] was used to construct and visualize the interaction network between genes and the transcription factor regulating them. Key term enrichment analysis for the genes corresponding to CpGs was done using DAVID [26].

## Results

### DNMTi treatment causes induction of methylation in a small portion of the CpGs

After quality control, we analyzed 369,886 CpGs across the genome of HCT116 cells from Yang *et al*. [15], and identified hypermethylation (Δβ ≥0.10) of 638 unique CpGs (0.02% of the total analyzed CpGs) in 393 unique genes after 5 days of decitabine treatment, as compared to untreated cells (Figure 1A). Most of them were hypomethylated in the untreated state (median β= 0.18), and after decitabine treatment, a median increase of 0.12 (Δβ = 0.12) in methylation level was observed for these sites. The detailed list of the identified CpGs is provided in Supplementary Table 1.

**Figure 1:**
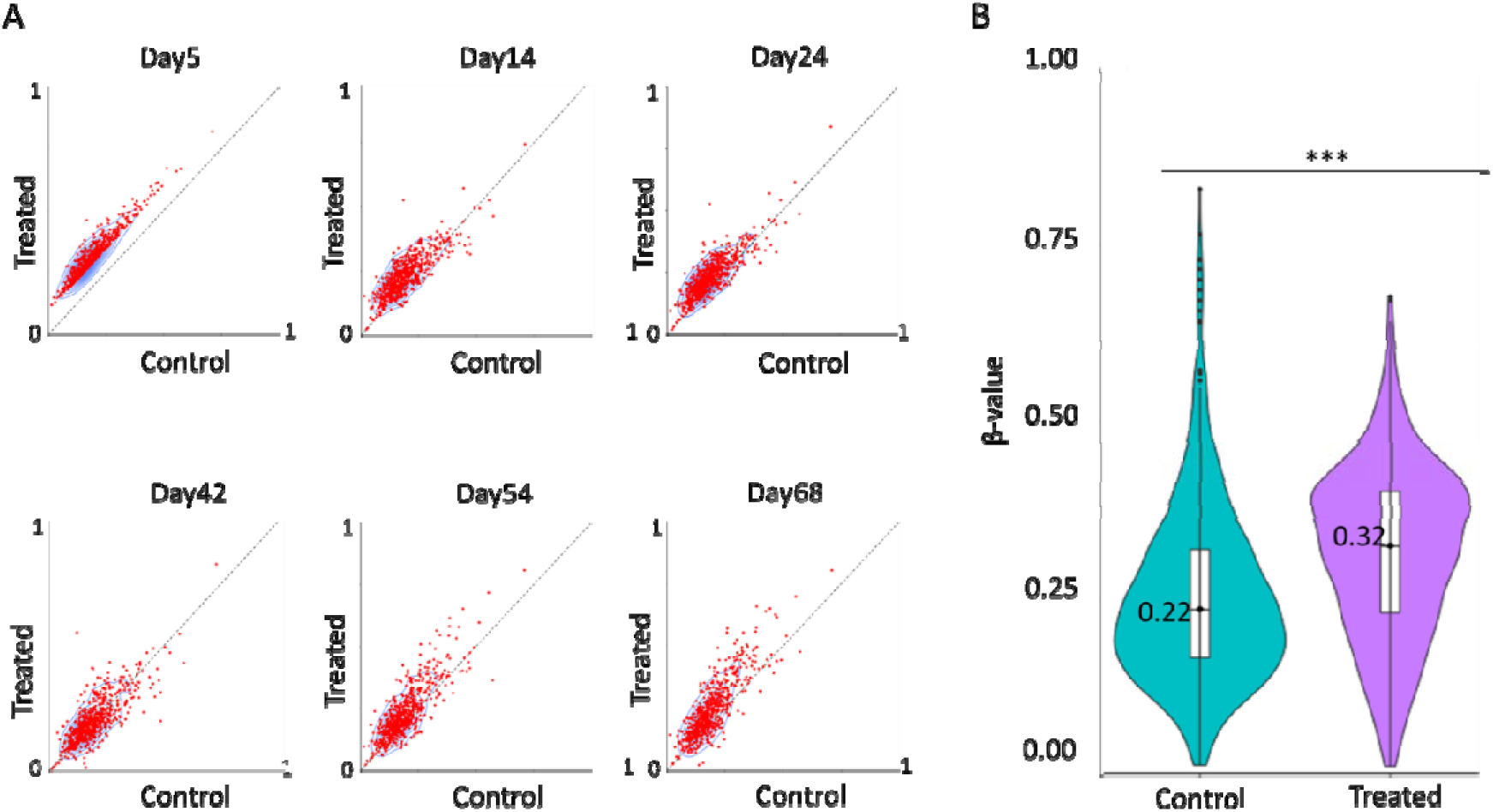
*decitabine treatment increases DNA methylation levels of a subset of CpGs. (A) Scatter plots showing DNA methylation patterns of 638 differentially methylated CpGs between untreated control cells and decitabine treated cells at various time points in* Yang *et al*. study [15]*. The x-axis indicates the DNA methylation level of probes in the untreated control, and the y-axis in decitabine treated cells. (B) Violin plot showing the median methylation level (horizontal line) and distribution patterns (density and IQR) of the identified 583 CpGs in untreated and decitabine treated HCT116 cells after 24 hours in* Han *et al* study [16]*. The statistical significance was assessed using the non-parametric Wilcoxon test. ***P<0.0005*.

Analysis of another methylation data for HCT116 cell line from the Han *et al* study [16] validated our finding, as we found a corresponding increase in methylation level (median increase in β values 0.09) after decitabine treatment (0.3 µM for 24 hours) at 583 (91%) differentially methylated CpGs that are common between the two studies (Figure 1B). These results indicate that the increase in DNA methylation of most of the identified sites starts as early as 24 hours after the DNMTi treatment and lasts up to at least day 5. The findings are also robust to common technical and processing variability across two laboratory conditions.

### Increase in methylation of identified CpGs is cancer type and tissue-specific

To test the effect of decitabine treatment on identified differentially methylated CpGs in other cancer cell lines, we re-analyzed publicly available data for decitabine-treated bladder cancer T24 cells. An increase in median DNA methylation levels (Δβ =0.14) at 616 (97%) common CpGs was observed after drug treatment (1 µM of decitabine for 24 hours) in T24 cells (Figure 2A). We further analyzed methylation level of identified loci for breast cancer cell line (MCF7), where cells have been treated with 0.06 µM of decitabine for 72 hours, but did not observe decrease in median methylation level (Δβ = -0.01) of 590 common differentially methylated CpGs (93%) in response to decitabine treatment (Figure 2B). These results indicate that methylation levels of the identified probes either increase after decitabine treatment in multiple cancer types or remain similar which is in contrast to the general effect of decitabine over CpGs methylation as we observed a significant decrease in the median methylation (Δβ = -0.14 for T24 and Δβ = -0.22 for MCF7) level of other CpGs present in the 450K beadchip in both of the cell lines (Supplementary Figure 1). Further, the result suggests that the degree of the change in methylation of these probes is highly cancer-specific.

**Figure 2:**
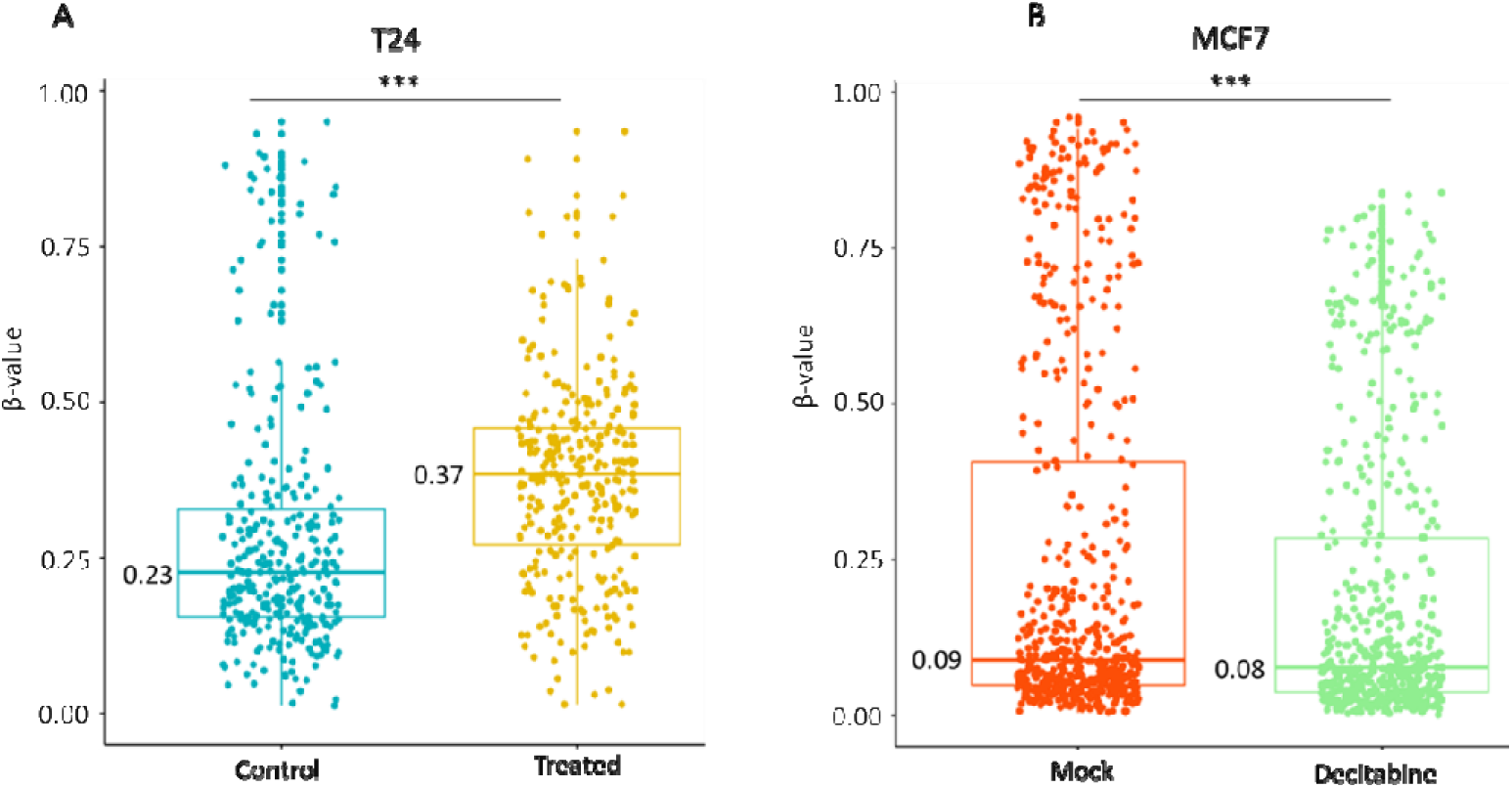
Increase in methylation of identified CpGs is tissue specific. (A) Methylation level of 616 identified probes in untreated and decitabine-treated bladder cancer T24 cell line after 24 hours of drug treatment. (B) Methylation level of 590 identified probes in mock (DMSO) treated and decitabine-treated breast cancer MCF7 cell line after 72 hours of drug treatment. The statistical significance was assessed using the non-parametric Wilcoxon test. ***P<0.0005.

### DNMTi treatment also increased methylation level of identified CpGs in colon, ovarian and breast cancer cells

To study the question whether the increase in methylation of the identified sites is decitabine-specific or whether also another DNMT inhibitor shows similar changes, we tested the methylation induction behavior of azacytidine, another FDA approved DNMT inhibitor, in multiple cancer cell lines. An analysis of 450K data from 52 cell lines revealed that azacytidine also increased methylation of identified CpGs in 9 out of 13 (69%) ovarian cancer cell lines(median Δβ >0.03), 3 out of 26 (11.5%) breast cancer cell lines(median Δβ >0.01), and in 9 out of 12 (75%) colon cancer cell lines (median Δβ >0.02), and 1 lymphoma cell line (U937, median Δβ >0.06) as shown in Figure 3A,B,C and D. Our analysis revealed that the increase in methylation level of identified loci in response to azacytidine treatment is not universal across all the cancer cell types, rather azacytidine treatment also causes an increase in median DNA methylation of identified sites in a tissue-specific manner.

**Figure 3:**
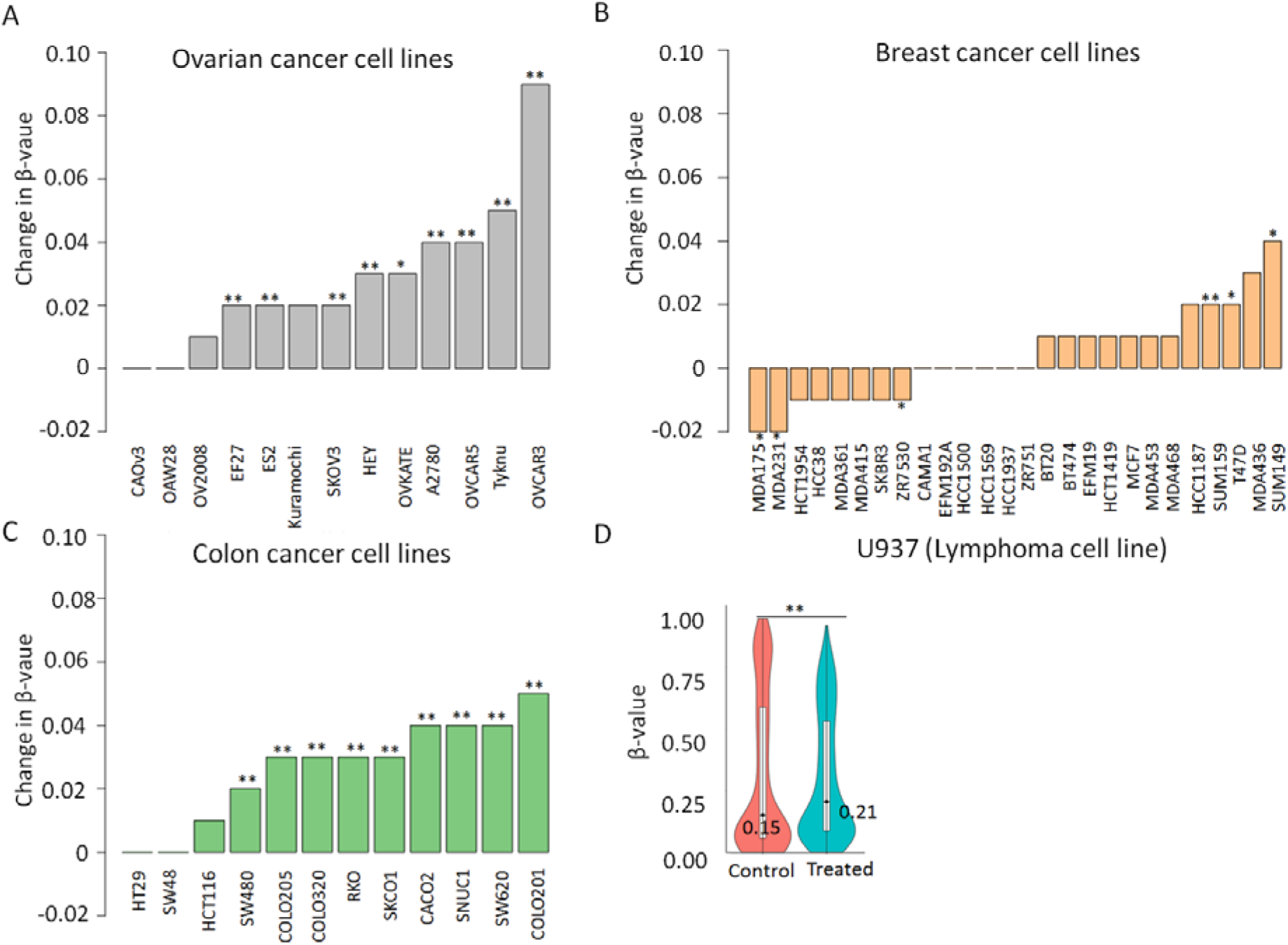
*Azacytidine treatment increases methylation of identified sites in a subset of cell lines. Change in median methylation level of identified CpGs in (A) 13 ovarian cancer cell lines (B) 26 breast cancer cell lines (C) and 12 colon cancer cell lines has been shown as barplot. The bar plot represents the difference in the median methylation level of identified CpGs between cells treated with 0. 5 µM azacytidine (test group) or carboplatin (mock group) after 72 hours. (D) Violin plot showing the distribution of methylation level of identified probes in untreated (control) and treated cells (*5mM for azacytidine for 72 hours) *in U937 lymphoma cell lines*. *The statistical significance was assessed using the non-parametric Wilcoxon test. *P<0.05, **P<0.005*

### Genes corresponding to the identified CpGs show differential expression in cancerous tissue

One of the common mechanism how DNA methylation affects the biological processes is by modulating the expression of the nearby genes. To test the functional role of the increase in methylation of the identified differentially methylated CpGs, we explored the correlation between DNA methylation and expression levels of the corresponding genes using data from HCT116 cell line after decitabine treatment at multiple time-points (0, 5, 14, 24, and 42 days). A strong correlation (|r|≥0.80) between the gene expression and DNA methylation profiles was observed at 26% (N =166) loci in HCT116 cells, out of them, 48 CpGs corresponding to 43 genes were significant (P<0.05). We observed a highly-significant correlation between the gene body CpGs (cg08099431) methylation in RORA gene and its expression level in HCT116 cells (r = 0.99, FDR = 0.01) in HCT116 (Supplementary Figure 2). The correlation plot for highly correlated CpGs (|r|≥0.80) falling in promoter and gene body regions in HCT116 cell line is shown separately in Figure 4A and 4D, respectively.

**Figure 4:**
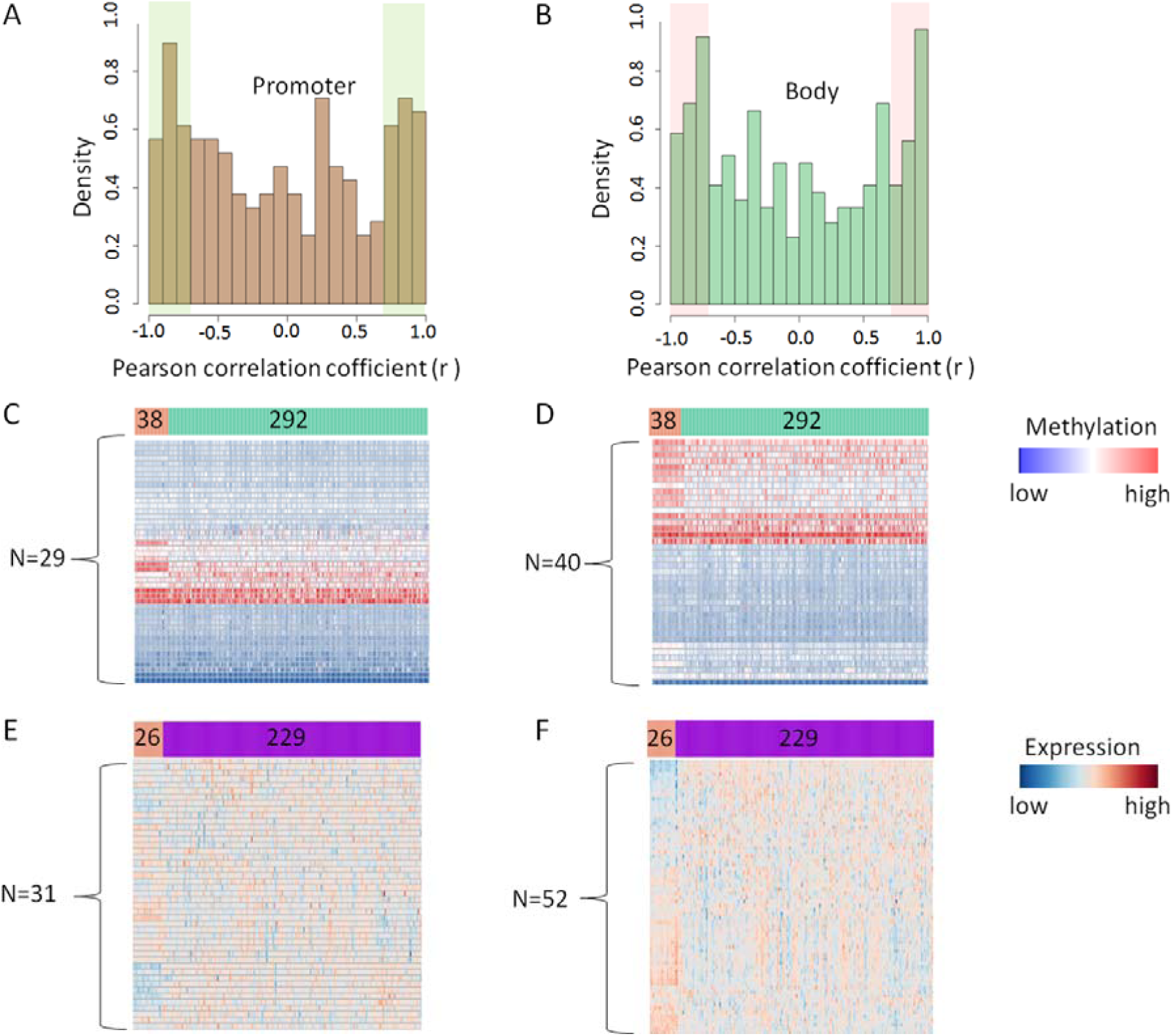
*DNA methylation and gene expression analysis of identified CpGs and showing a strong correlation* (|r|≥0.80) *between expression and methylation in HCTT116 cell line in TCGA colon adenocarcinoma samples.* (A) Distribution of correlation between DNA methylation and expression at the promoter region (left panel) and gene body (right panel) in HCT116 cell line. The x-axis denotes the Pearson correlation coefficient between gene expression and methylation in HCT116 cell lines and the y-axis the kernel density (B) Heatmap showing methylation level of the identified CpGs in TCGA colon samples. Forty-two out of 53 CpGs were available for promoter region (left panel), while 62 out of 86 CpGs were available for gene body (right panel) in TCGA datasets. Salmon color represents normal colon tissues (n=38); cyan color represents colon tumors (n=292). Blue indicates low beta values; red represents high beta values. (C) Supervised clustering of TCGA colon samples based on expression level of genes corresponding to differentially methylated CpGs. Separate heatmap showing expression level of 45 genes corresponding to the sites in promoter region (left panel), and 67 genes in the gene body region (right panel). Blue indicates low expression level; red represents high expression level.

To investigate the clinical relevance of increase in methylation of identified CpGs, we analyzed the methylation level of the 109 common CpGs showing strong expression-methylation correlation (|r|≥0.80) in HCT116 cell line in TCGA colon adenocarcinoma samples. This analysis revealed that 43% (47 out of 109) of the correlated CpGs were also differentially methylated between healthy and cancerous colon tissues in the TCGA data (FDR <0.05, Figure 4B and E, Supplementary Table 2). Differential expression analysis revealed that 77% of the corresponding genes (N=83 out of 112 genes for which data was available in TCGA) were also differentially expressed between the colon and normal tissues (FDR <0.05, Figure 4C, Supplementary Table 3). The differential expression of genes corresponding to the identified CpGs in cancerous tissues indicates that the increase in their DNA methylation is pathological in colon cancer.

### Genes with increased methylation are enriched in cancer-related pathways and are NFAT, LEF1, MAZ-regulated

We next investigated the functions of the genes corresponding to the identified CpGs using the GeneCodis (v2) gene set enrichment analysis tool. Gene ontology enrichment analysis revealed that five out of the 10 (50%) most significant GO processes (FDR = 0.05) were related to transcription regulation, which is one of the key functional role of DNA methylation in order to control gene expression (Figure 5A). Notably, the list of enriched genes included well-known oncogenes, such as AFF3, CTNND2, ELK4, ESR1, PAX3, TRRAP, and WHSC1L1. The pathway enrichment analysis revealed that olfactory transduction and p53 signaling pathway were overrepresented (FDR= 0.05) in the gene set with increased methylation (Figure 5B). Enrichment analysis further revealed that the corresponding genes were enriched for alternating splicing as the major keyword (fold enrichment = 1.27, FDR = 0.000132, in Figure 5C).

**Figure 5:**
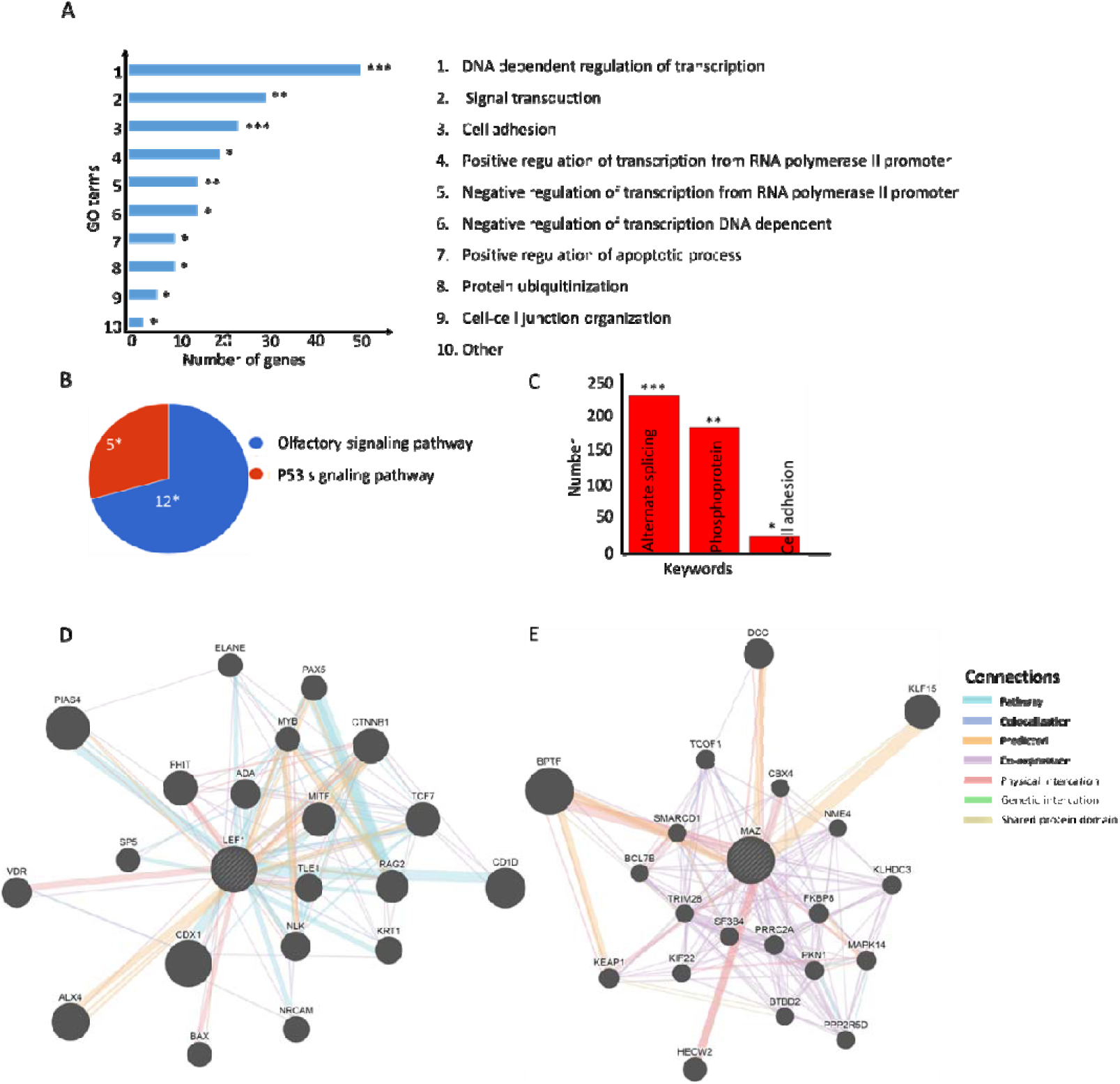
Gene ontology (GO) and pathway enrichment analyses of genes corresponding to differentially methylated CpGs. (A) Top ten significantly enriched cellular processes have been shown as bar plot. The lengths of the bars denote the number of genes present in each of the top GO categories. (B) Pie-chart showing the significantly enriched pathways for the genes. The number of genes present in each pathway group has been shown along with the hypergeometric test p-value corrected for multiple testing. (C) The keywords enrichment analysis for the genes has been shown as bar chart. The length of the bar represents the number of genes enriched for each keyword, the FDR-corrected p-value has been shown at the top. (D) Interaction network of genes regulated by LEF1 (E) Interaction network of genes regulated by MAZ. Only interactions among those genes that are directly connected to enriched transcription factors (LEF1 and MAZ) have been shown as network using GENEMANIA. The size of the nodes is proportional to score calculated by GENEMANIA using label propagation algorithm that indicates the relevance of each gene to the original list of genes based on the selected networks. *P<0.05, **P<0.005, ***P<0.0005

Further, enrichment analysis among the transcription factors regulating these identified genes revealed that 47 (12%) of genes were regulated by the nuclear factor of activated T-cells (NFAT, P =1.48x10^-8^, Supplementary Figure 3), 59 (15%) genes were regulated by lymphoid enhancer-binding factor 1 (LEF1) (P = 2.1 × 10^−8^, Figure 5D), and 51 (13%) genes by MYC Associated Zinc Finger Protein (MAZ) (P = 4.07× 10^−8^, Figure 5E). Therefore, by increasing DNA methylation, decitabine strongly affected several well-known oncogenes related to cancer-related pathways, especially the olfactory pathway and p53 tumor suppressor pathway, and a majority of these genes were regulated by NFAT, LEF1 and MAZ transcription factor (Table 1).

**Table 1:**
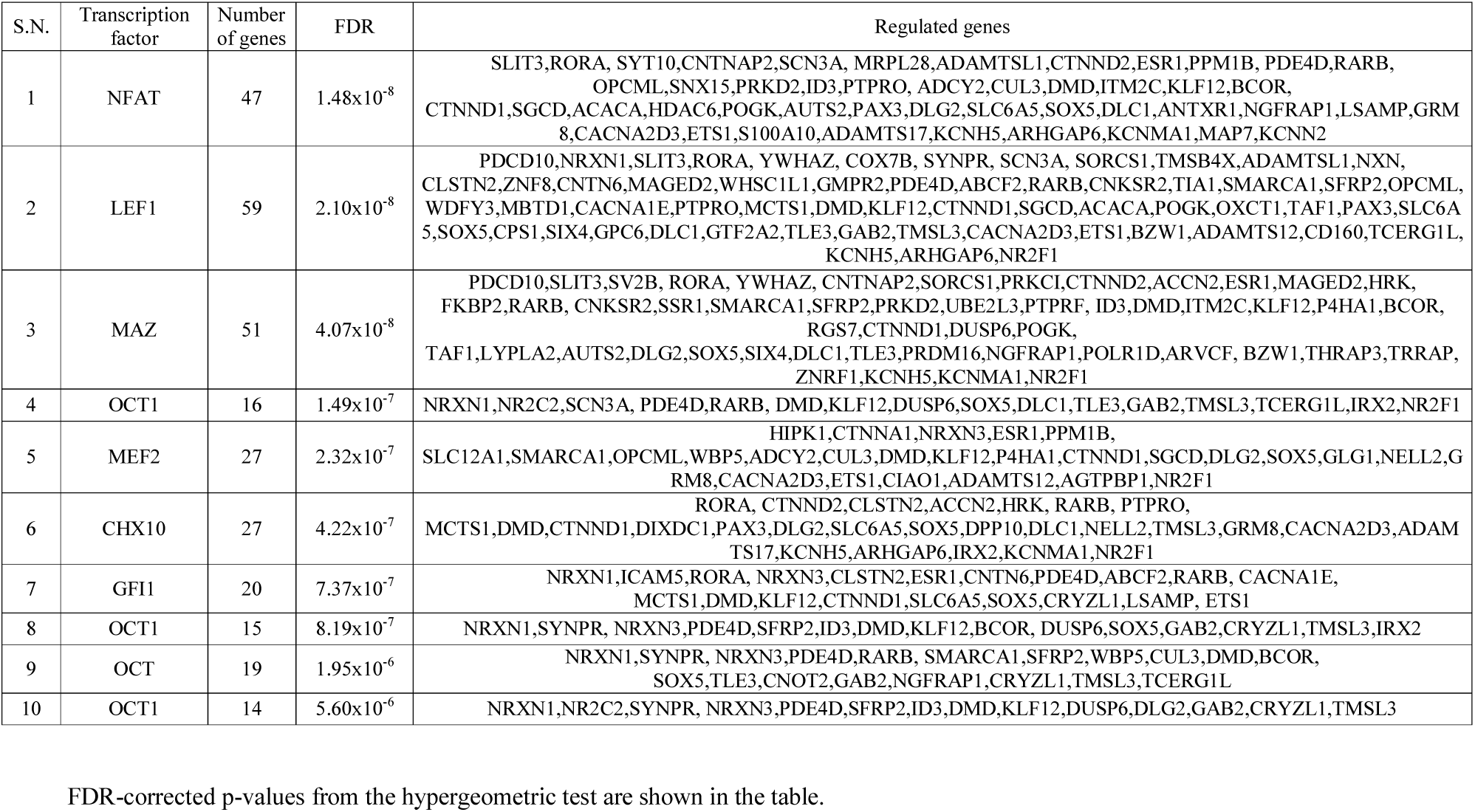
Top 10 transcription factor enriched in gene set corresponding to induced CpGs and its regulated genes

## Discussion

Our study indicates that clinically feasible dose of decitabine (0.06 µM to 300 µM) treatment causes a transient increase in DNA methylation level of a small fraction of CpGs in the genome related to critical signaling pathways involved in tumorigenesis. The use of 3-day exposure with such doses *in vitro* produces a quick increase in DNA methylation that may reflect the immediate response of cells to external stimuli. However, increase in DNA methylation in most of the sites were not correlated with the corresponding gene expression levels, instead, they were enriched for alternative splicing as the key process term (Supplementary figure 4). Previous results suggest a complex nature of DNMTi action over cancer cells by not only changing the expression of certain genes, but also regulating the number of different transcript isoforms for several other genes [27]. One of the major role of DNA methylation is to regulate alternative splicing in the genome mainly by modulation of the elongation rate of RNA polymerase II (Pol II) by CCCTC-binding factor (CTCF) and methy-l-CpG binding protein 2 (MeCP2) [28]. Increase in DNA methylation can also enhance alternating splicing by the formation of a protein bridge by heterochromatin protein 1 (HP1) that recruits splicing factors onto transcribed alternative exons [29].

We believe that the transient increase in methylation due to decitabine treatment is mainly for the temporary alteration of transcript level of certain genes rather than permanent shut down or enhancement of expression as the increase in methylation is transient (vanishes 10 days after treatment). However, further study is needed to understand how the induced methylation due to DNMTi treatment affects alternating splicing in cancer cells.

Further, genes corresponding to identified CpGs with increased methylation were enriched among cancer-related pathways (olfactory and p53 pathway), and are regulated by Nfat, Lef1 and Maz transcription factor. Lef1 is a known target gene of Wnt/β-catenin pathway and is upregulated in colonic carcinogenesis where Wnt-3A/beta-catenin signaling induces transcription from the LEF-1 promoter [30]. Knockdown of LEF1 inhibits colon cancer progression *in vitro* and *in vivo* [31]. Further, enriched p53 pathway and olfactory receptor signaling pathways are a hallmark of multiple cancer type, and are involved in cell proliferation, migration, and apoptosis of cancerous cells [32, 33, 34]. Olfactory receptor and related signaling activation inhibit cell proliferation and apoptosis in colorectal cancer cells [33]. Enrichment of genes among the cancer-related pathways involved in cell proliferation, differentiation, and apoptosis suggests a non-random, systematic selection of genes for a transient increase in methylation in order to carry out cancer-related biological process in cells.

Based on our analysis, there are multiple reasons to suggest that at least one key mechanism underlying the anti-tumor responses to DNMTi treatment may involve an increase in DNA methylation level of specific genes in cancer cells. First, we showed an increase in the pattern of DNA methylation in more than one type of cancer cells. Second, as defined for an epigenetic change, these sustained changes persist for significant periods of time (at least more than 5 days) after a transient, subsequently withdrawn, drug exposure (in this case 72 hours). Third, the expression patterns for a subset of the genes are different between cancer and normal tissue types. Importantly, these changes are induced by drug doses that do not acutely kill cells and, thus, allow the transient alterations in gene methylation patterns to act on emerging molecular changes to cells after DNMTi therapy. We showed that these changes include anti-tumor events in multiple key pathways, such as p53 pathway, olfactory receptor pathway, regulation of transcription level, and others which can cause huge molecular cascade in the cells even after removal of the drug. Thus, increased methylation might be considered as a key feature of DNMTi therapy that can alter multiple cancer-related pathways simultaneously.

## Conclusion

In summary, our findings provide novel insights into understanding the mechanism of action of DNMTi treatment in case of multiple cancer types, primarily colon cancer. Our findings suggest the existence of CpG sites in the genome that can resist DNMTi treatment and show an opposite effect of hypermethylation than expected demethylation, hence these CpGs could be clinically applicable as a response-predictive biomarker for patient stratification. Our results also suggest that DNMTi has a complex mechanism of action and a generalized pattern for the activity of DNMTi is challenging to find. Hence, the effects of DNMTi on cancer tissues should be analyzed at the individual gene level, rather than at the entire genomic level, and separately for each tissue type and even cancer patient.

### Future perspective

Our analysis is the first of its kind that directly shows increased methylation level at certain loci after DNMTi treatment in the HCT116 genome, contrary to its classical well-studied role in decreasing methylation. These findings were also validated across multiple cells lines belonging to different cancer and tissue types. However, a more functional mechanistic study in higher model systems and human tissue types is required for revealing how the increase in methylation at individual loci alters treatment response and the pathological burden of disease. We hope that these current observations will have implications for further research about DNMTi response, as well as resistance mechanisms, with the aim use drug-induced methylation with DNMT inhibitors as tool for treatment strategy for multiple cancers.

### Summary points

- DNMTi treatment increases DNA methylation in a small fraction of loci in HCT116 cells.
- The increase in methylation is transient and exists between 24 hours to at least 5 days.
- There is a limited correlation between DNA methylation of CpGs with increased methylation and gene expression and most of the genes with such CpGs are enriched for alternating splicing.
- A subset of the CpGs with increased methylation is differentially expressed between cancer and healthy tissue in the TCGA colon cancer data.
- 77% of genes having CpGs with increased methylations showed differential expression between the colon and normal tissues.
- Identified CpGs sites are enriched for enhancer regions in the genome.
- Genes corresponding to differentially methylated CpGs are enriched for p53 and olfactory receptor signaling pathways and are involved in transcriptional regulation of cells.
- These data suggest a complex nature of decitabine action on the genome and its effect need to be analyzed in a specific genetic context, instead of using pan-genome analysis.

---

**Supplementary Table 1:**
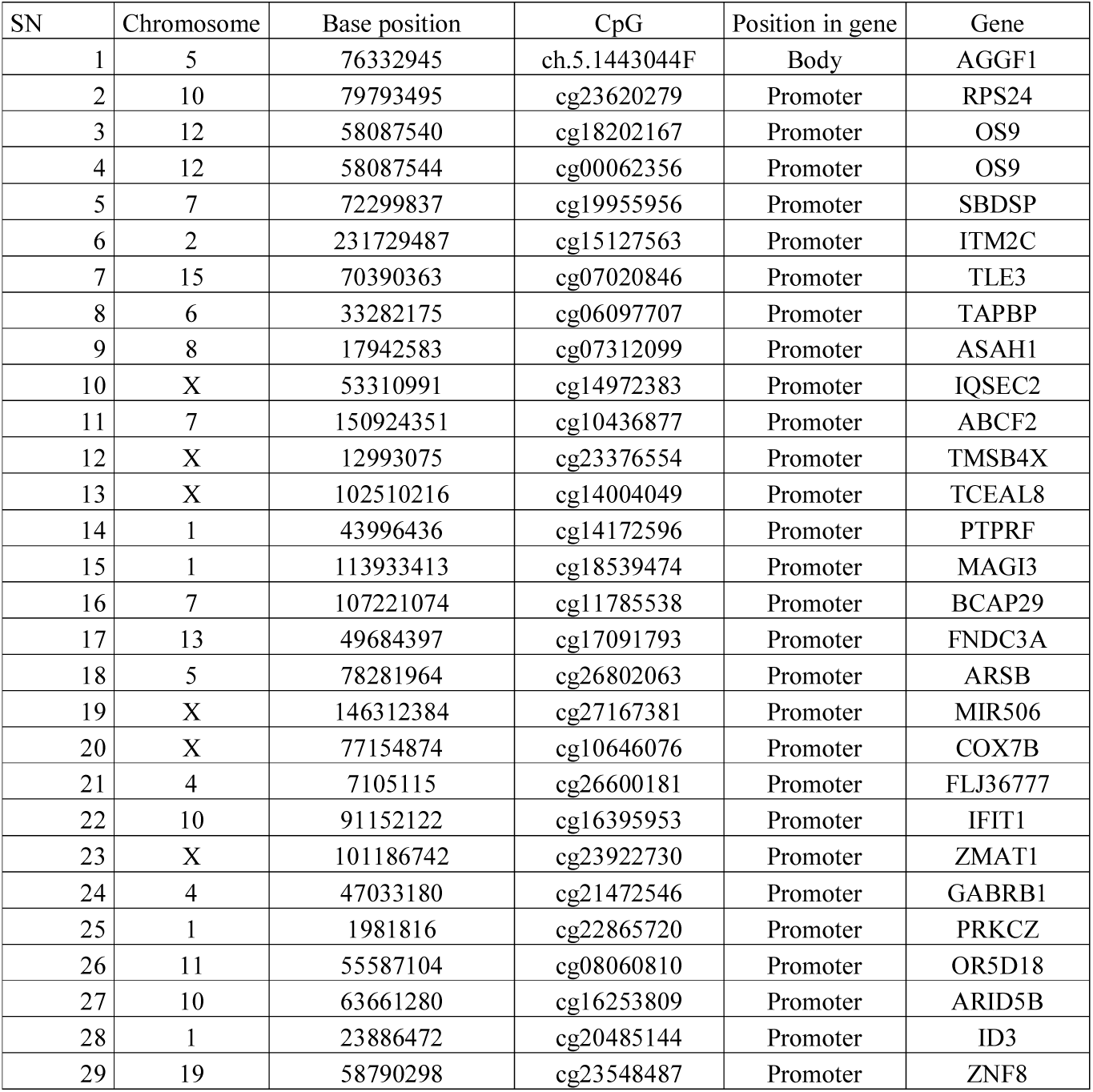
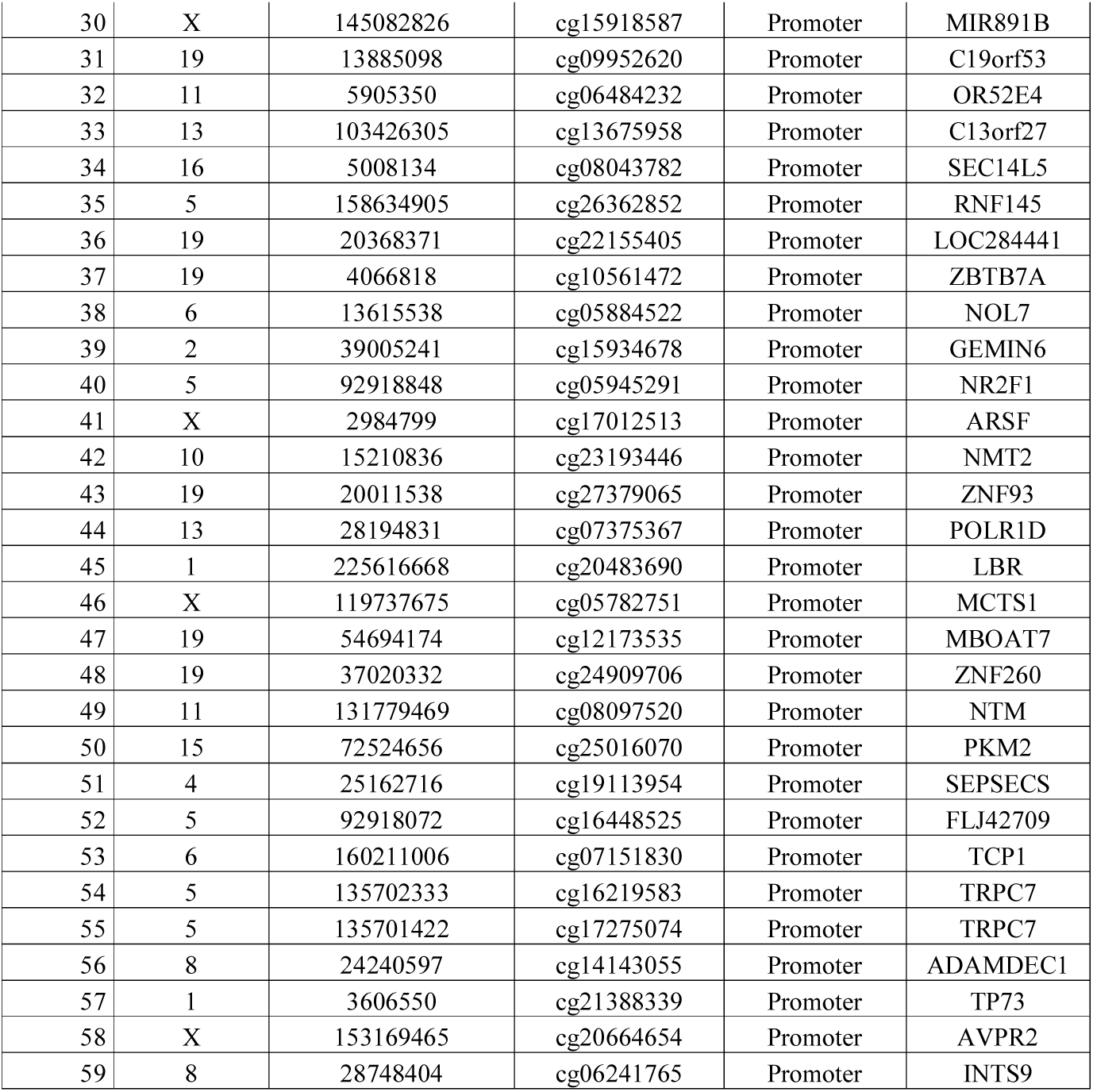
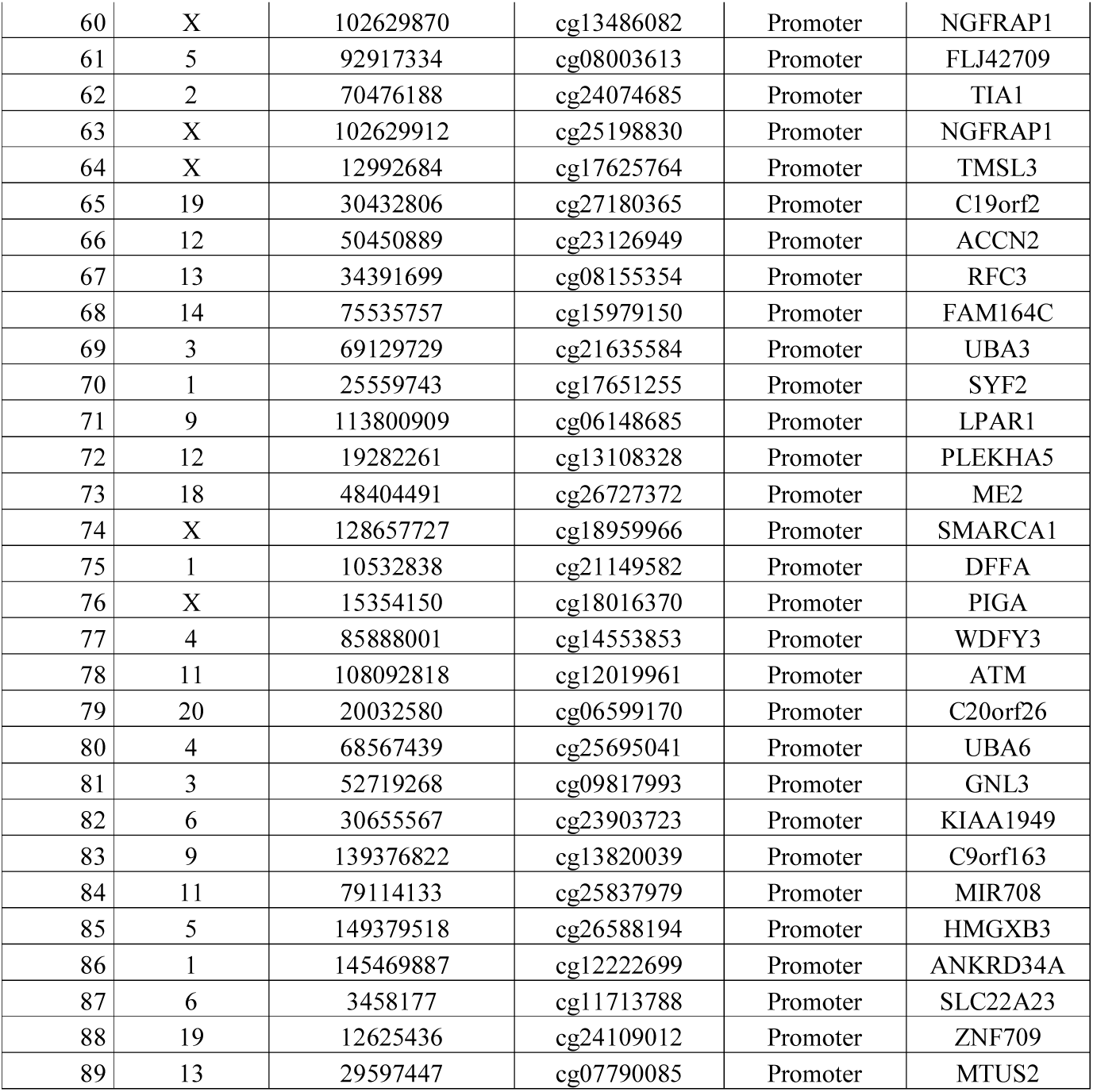
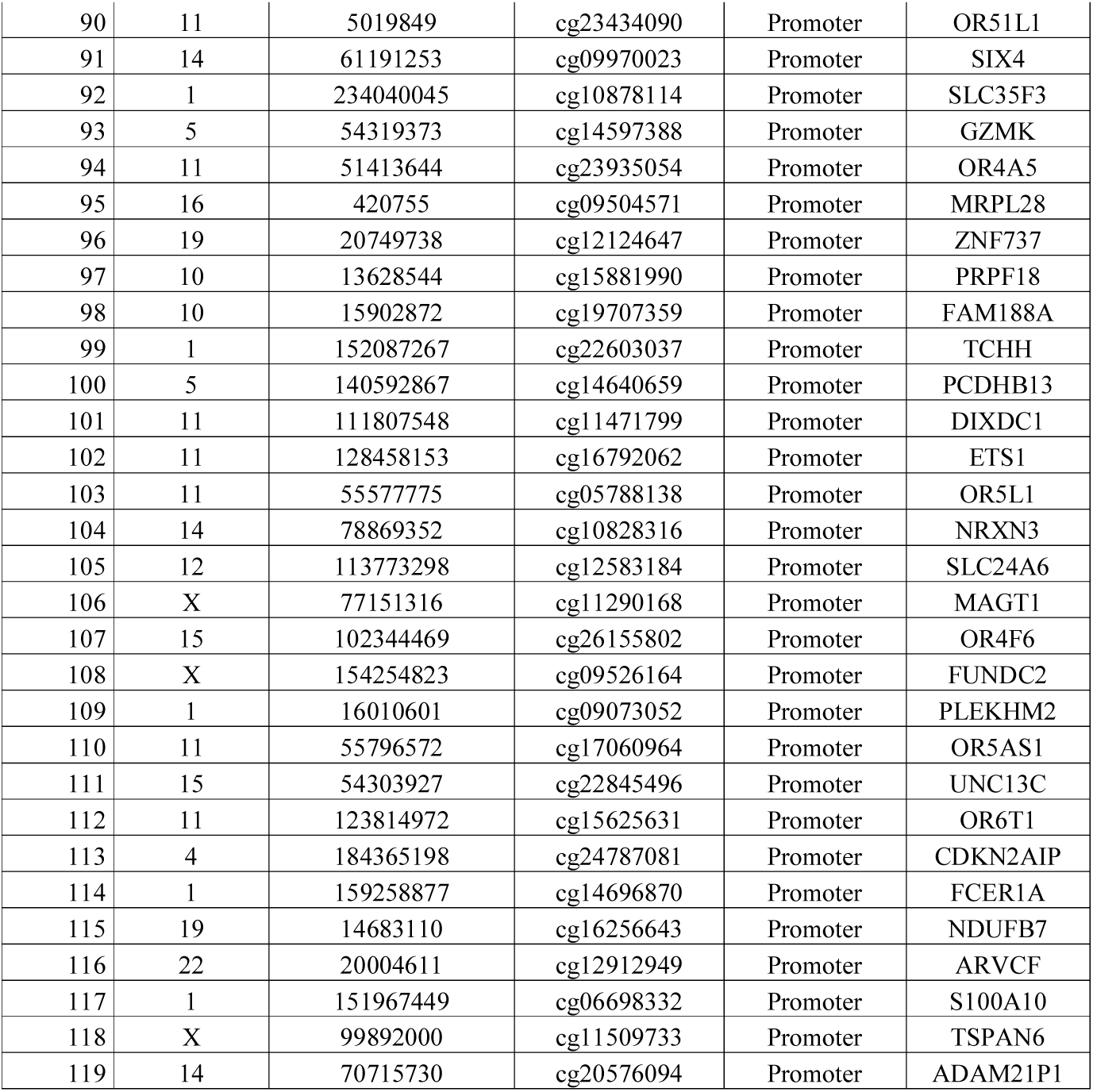
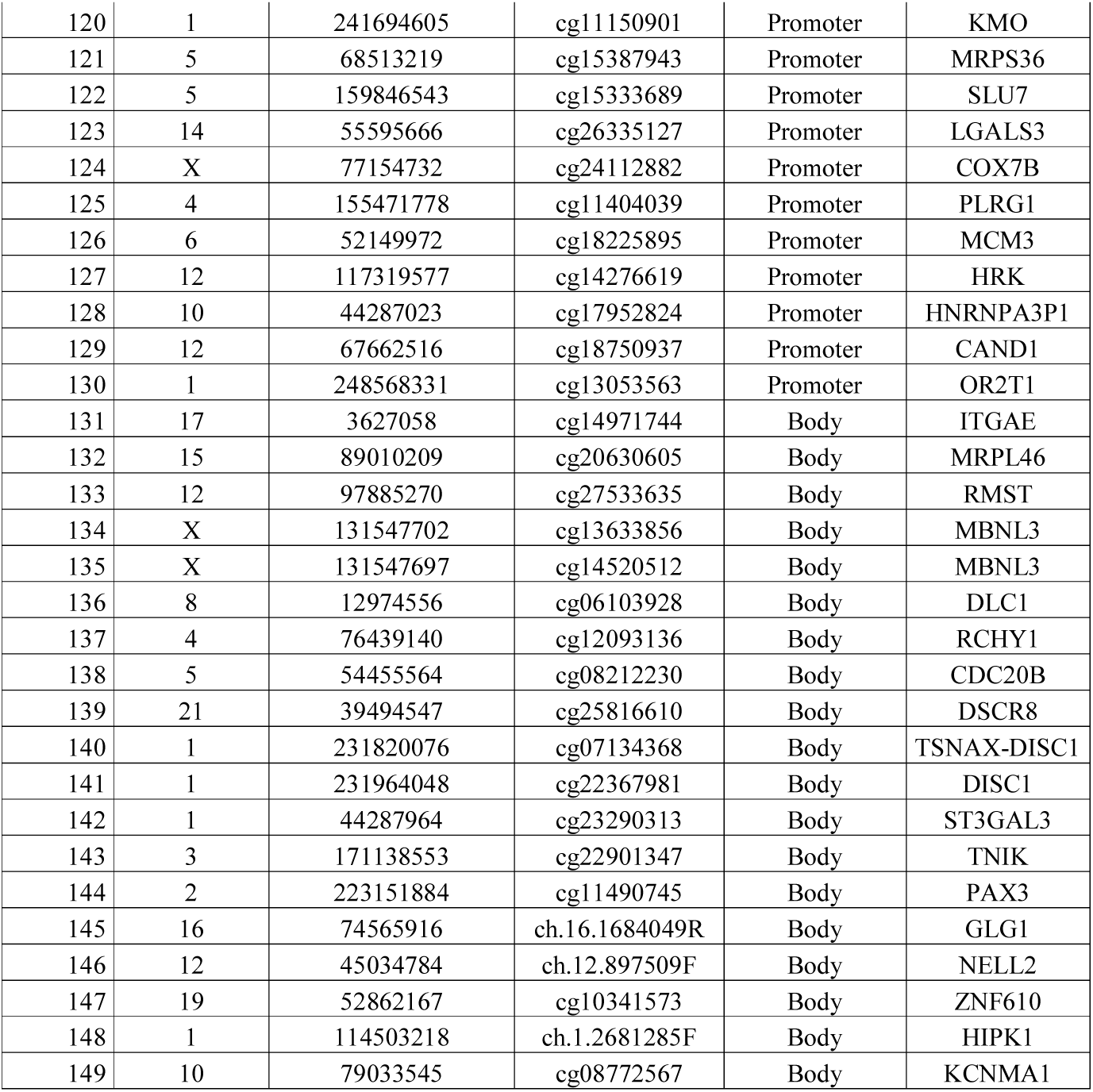
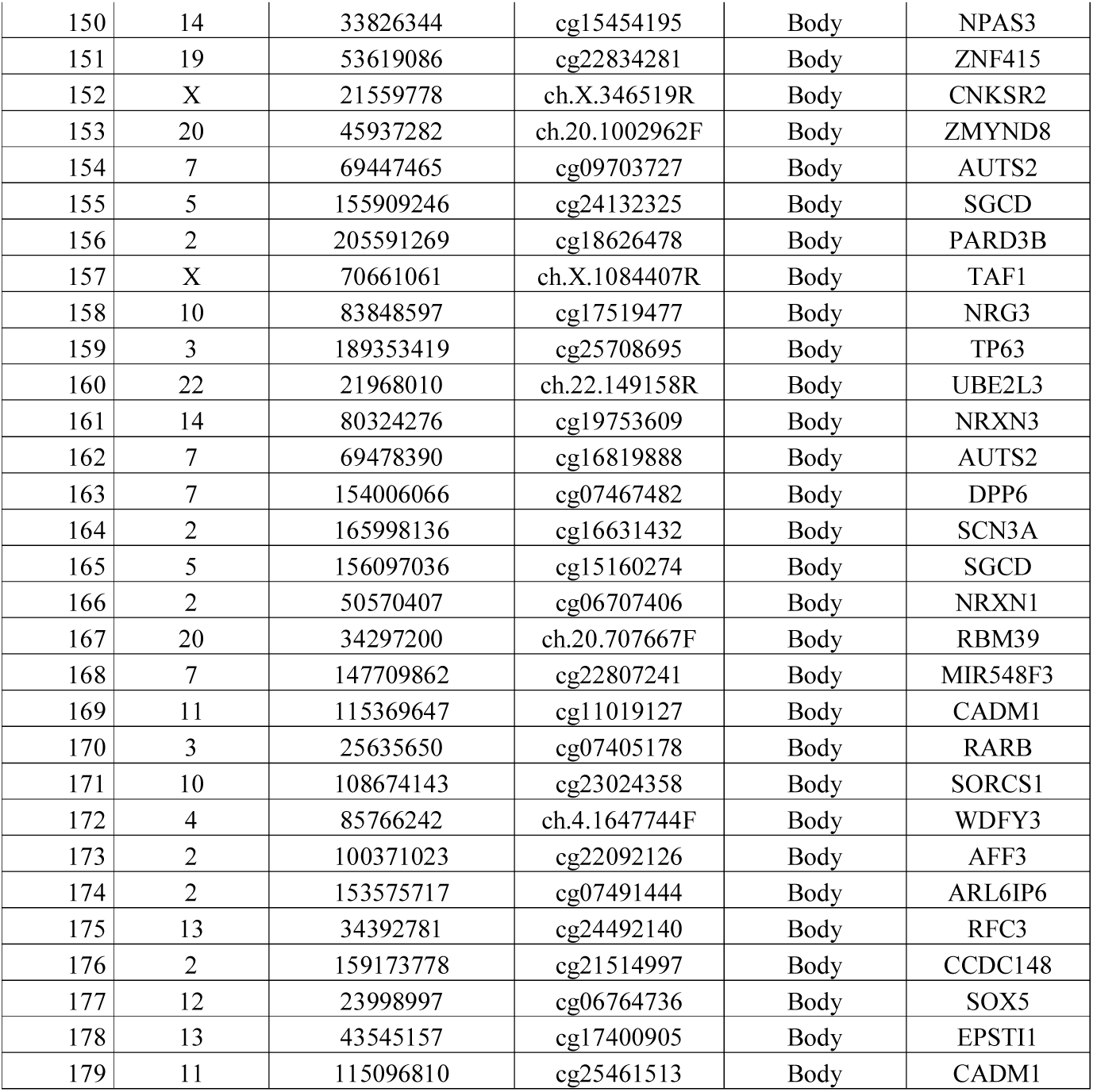
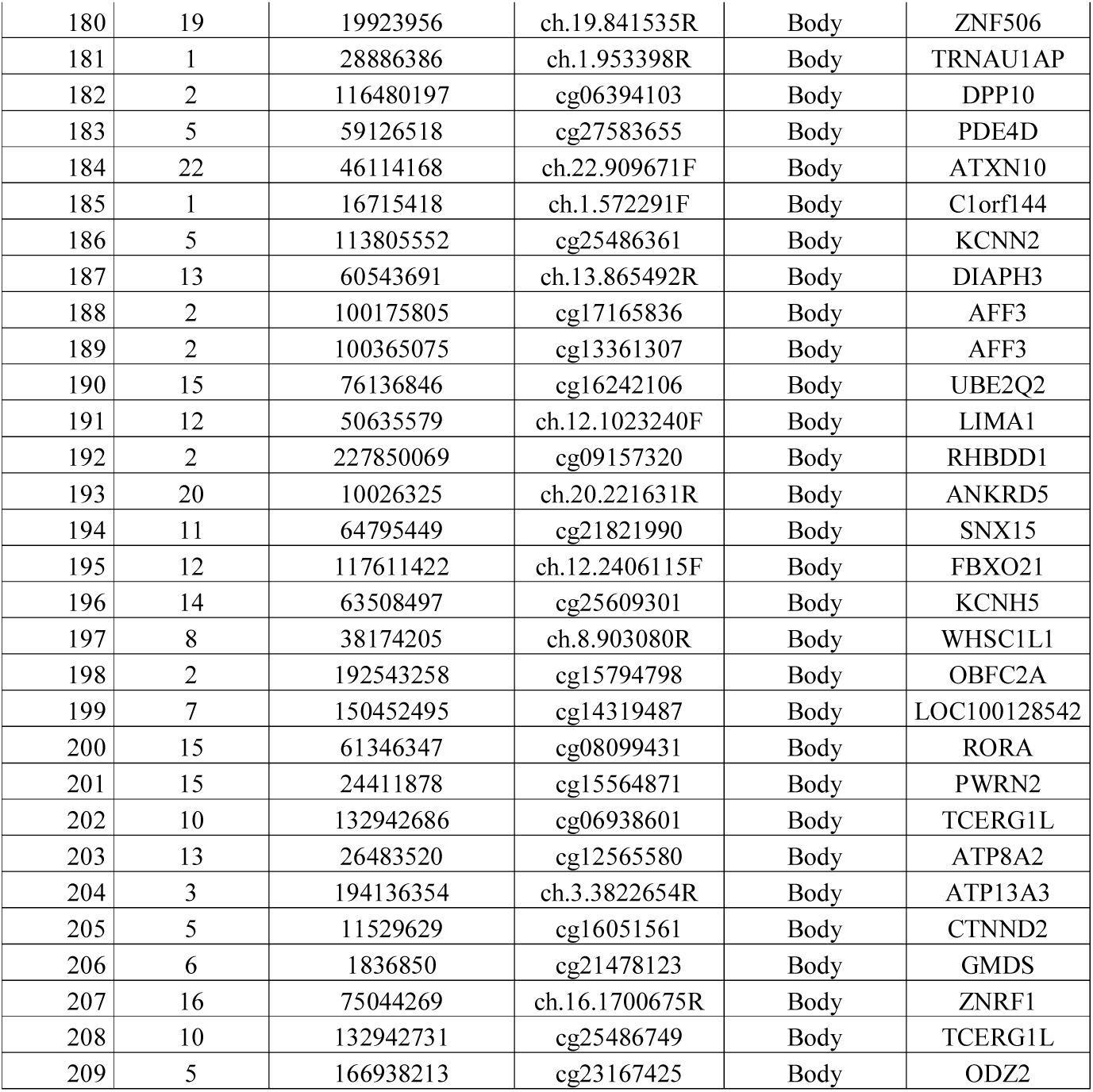
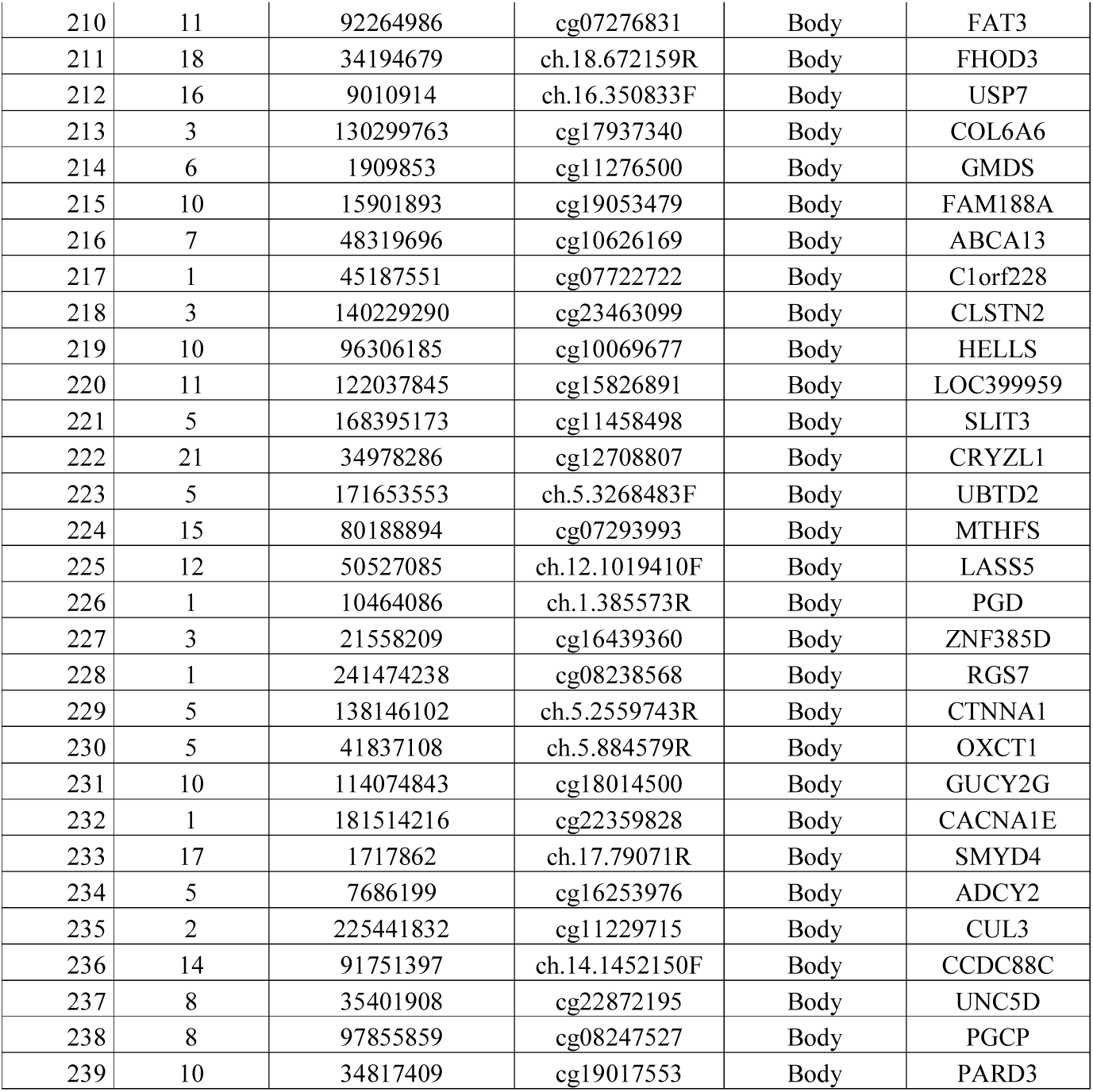
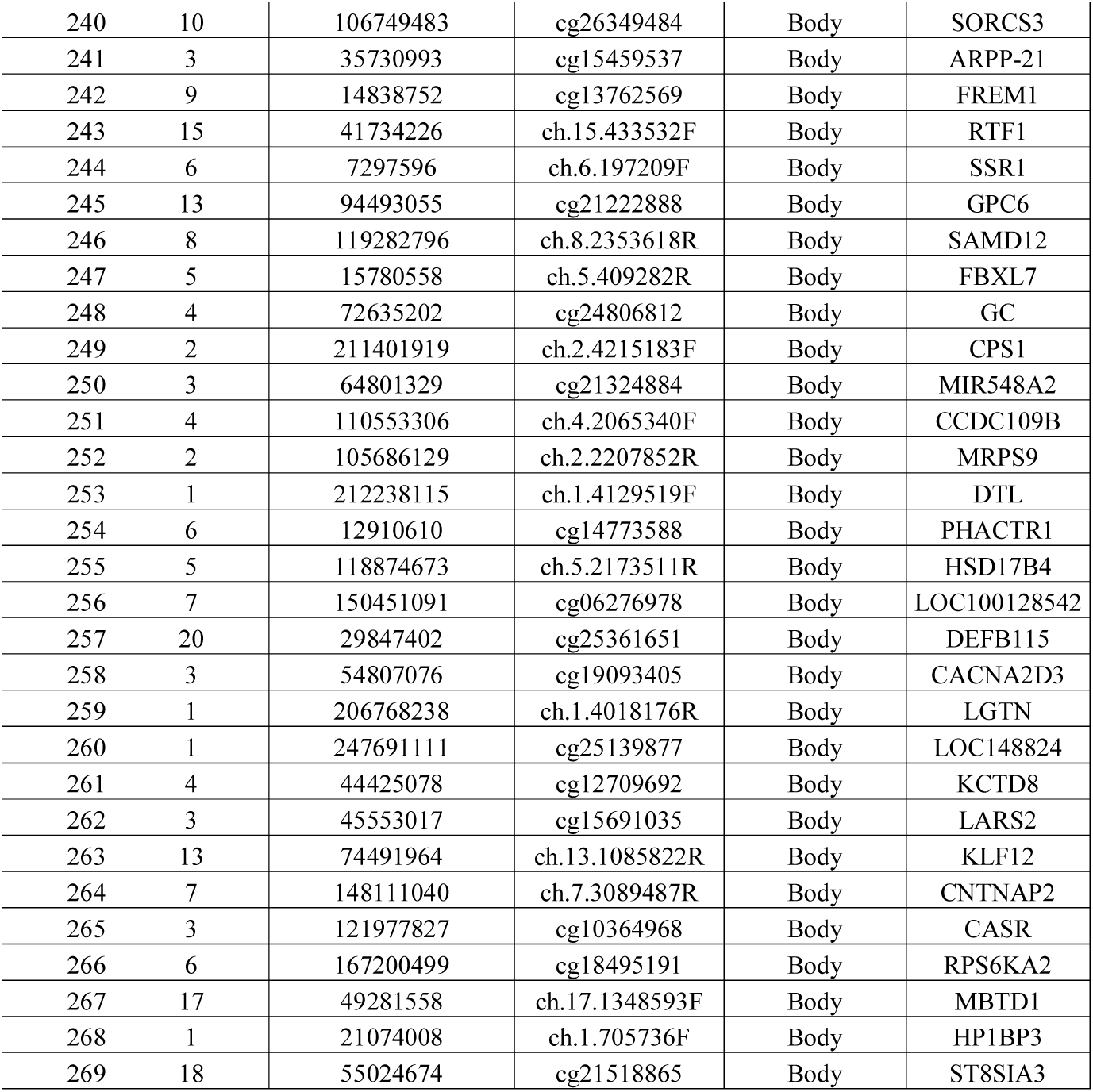
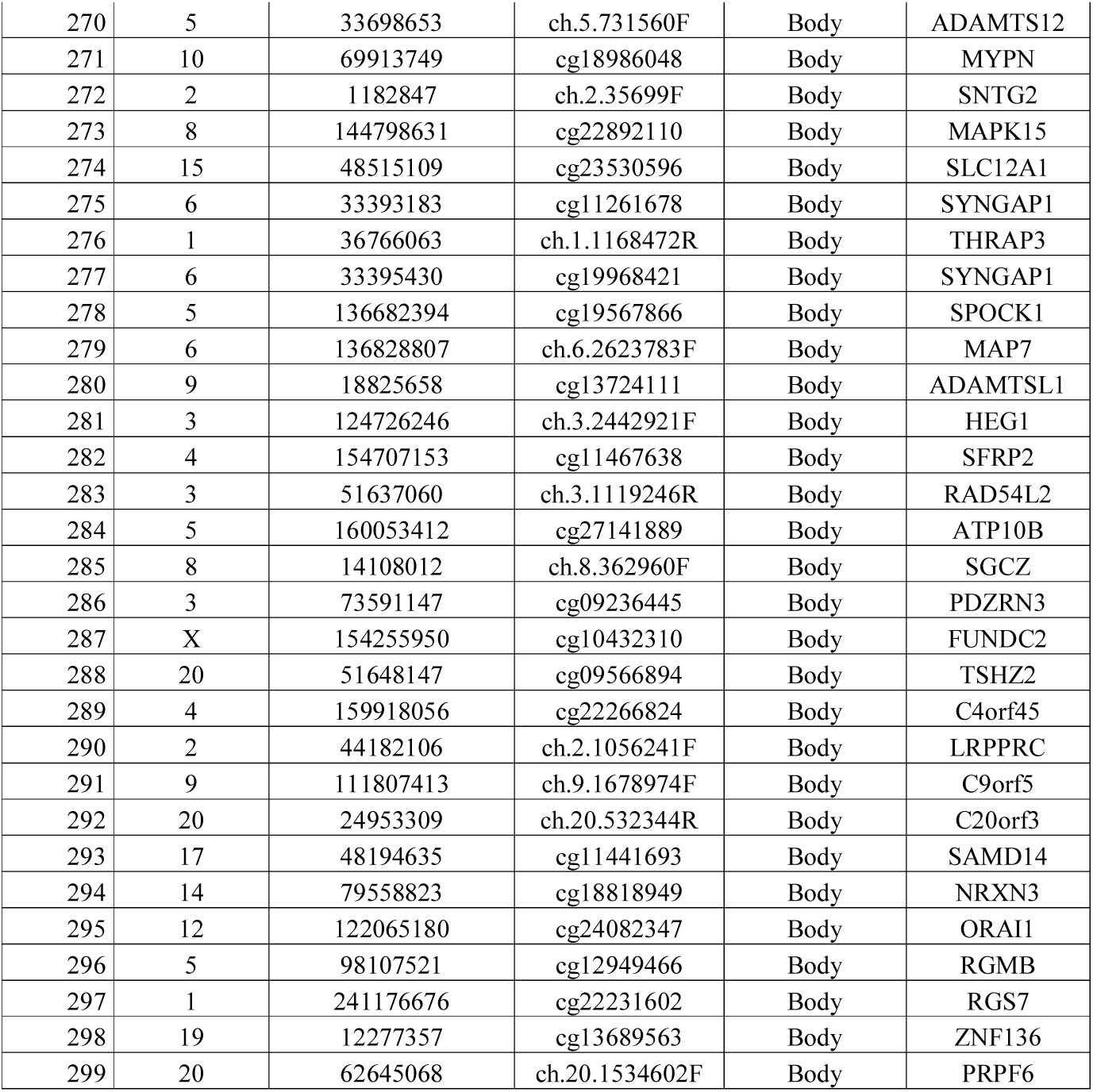
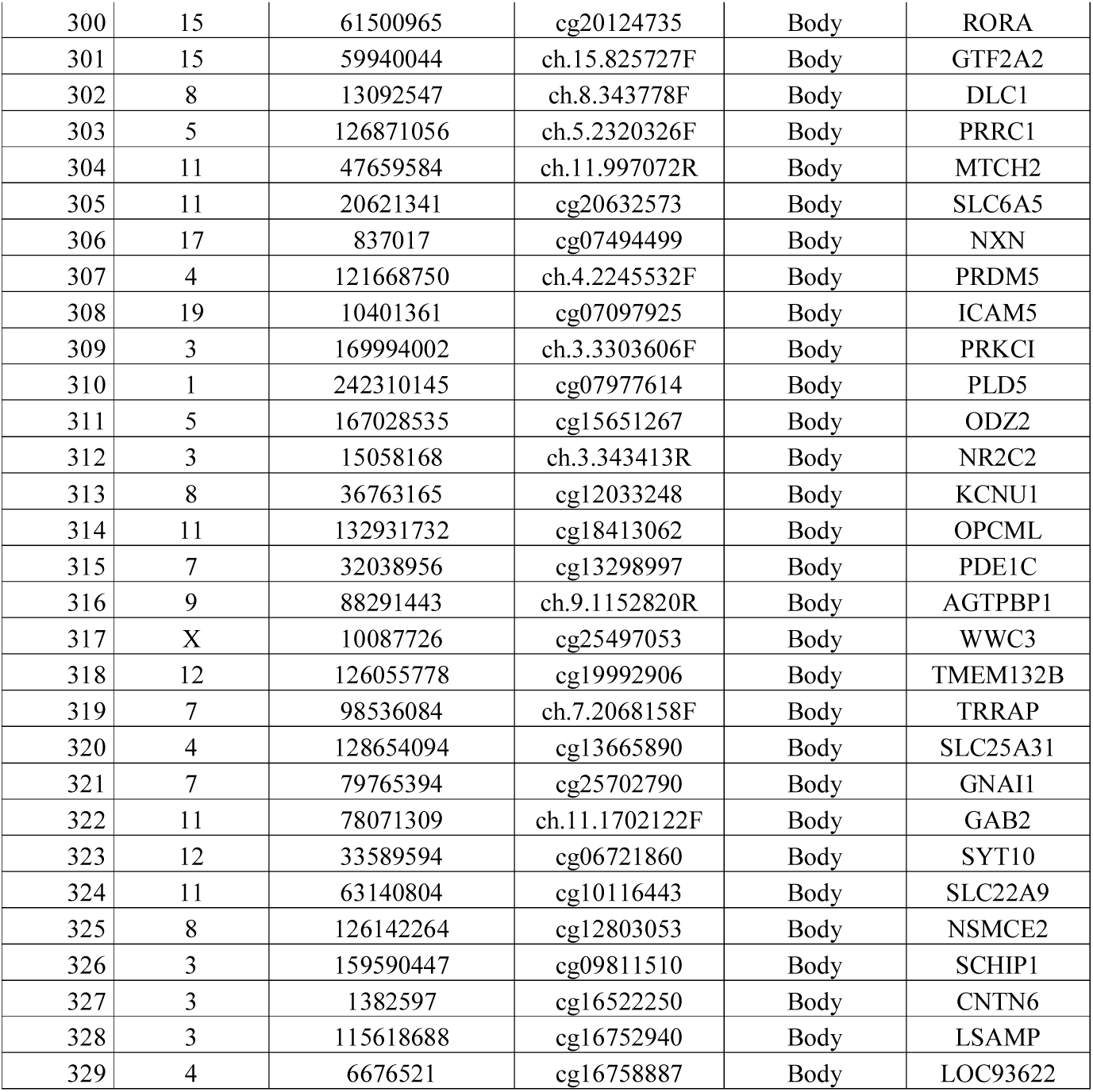
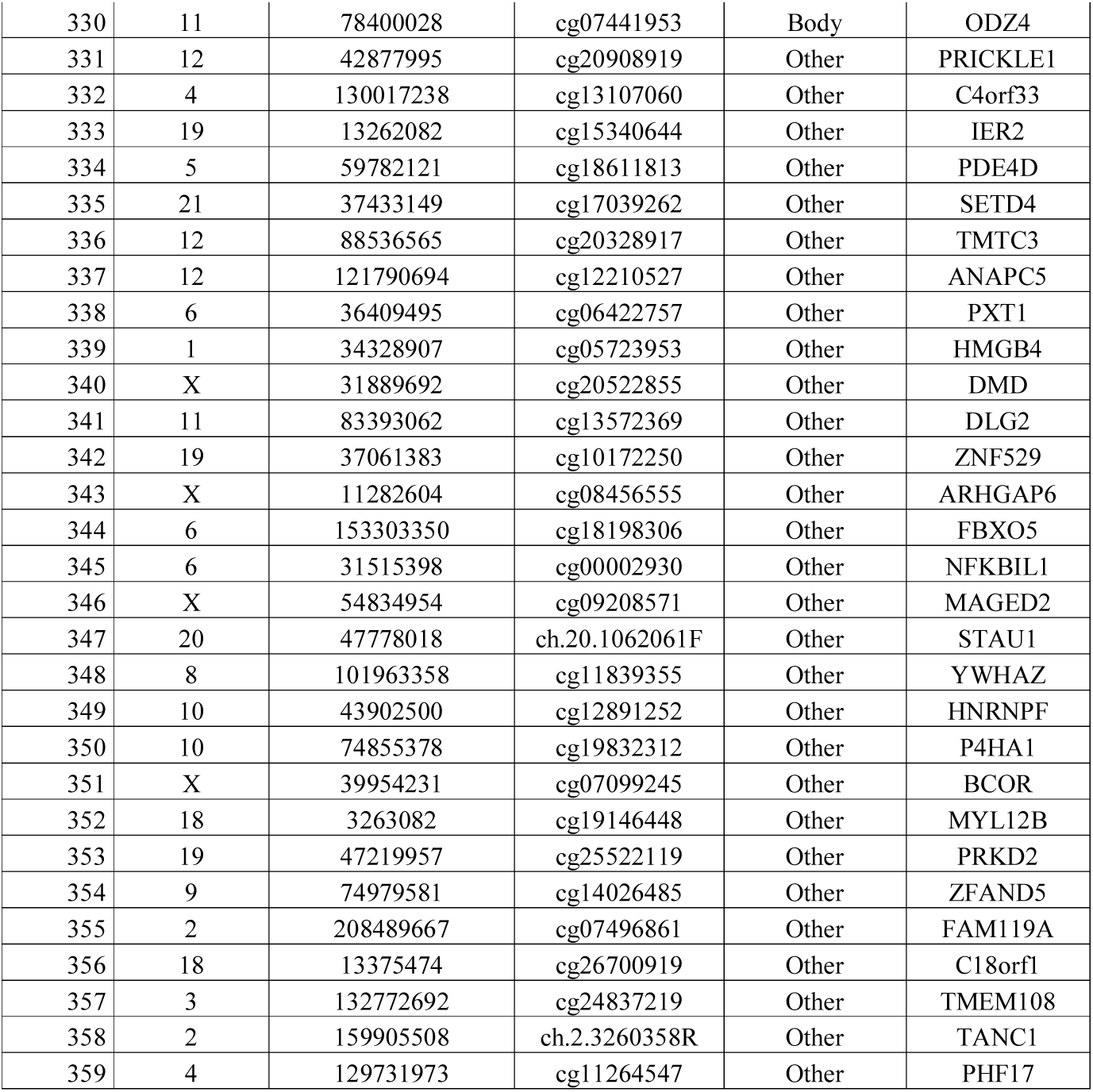
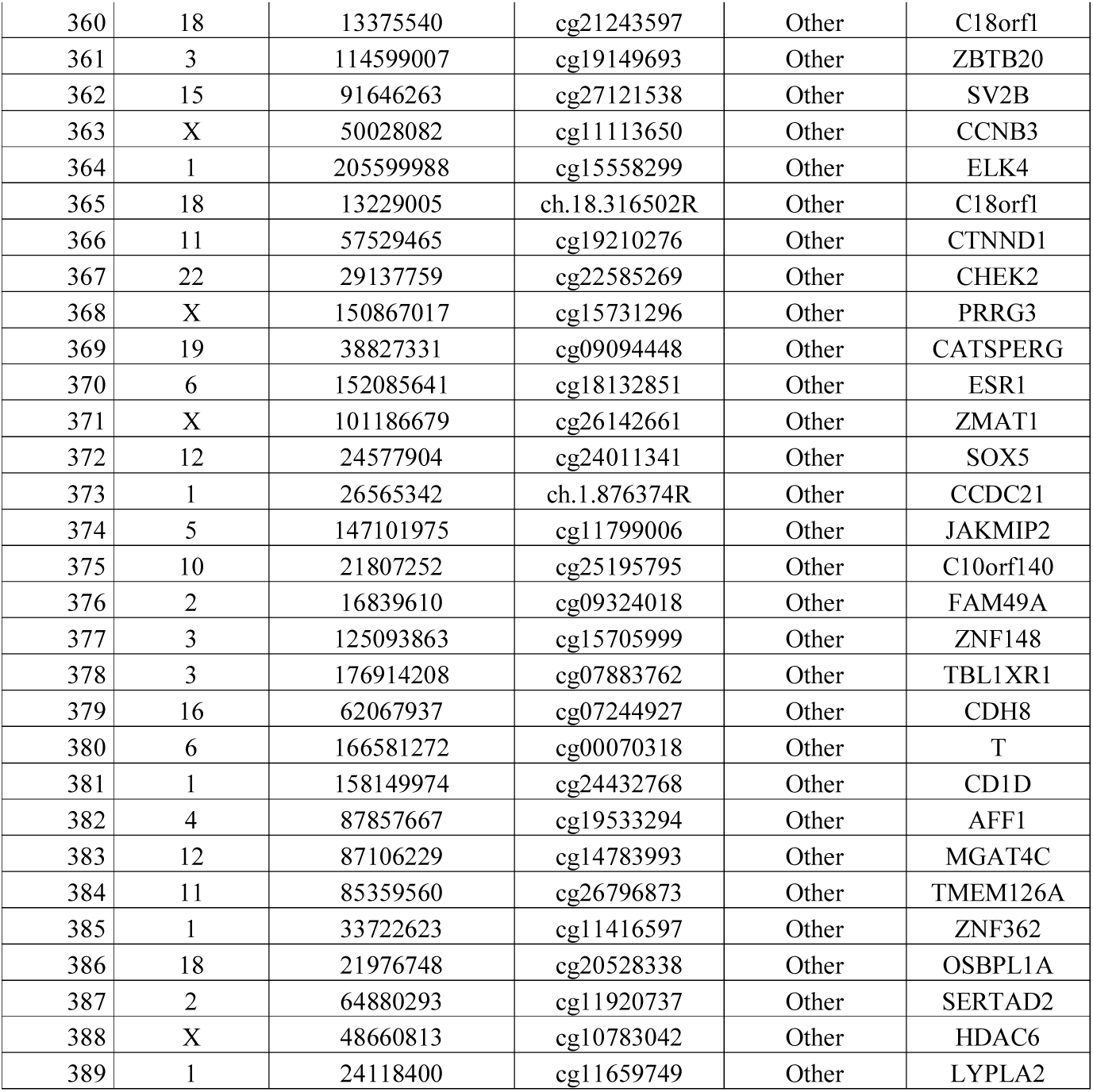
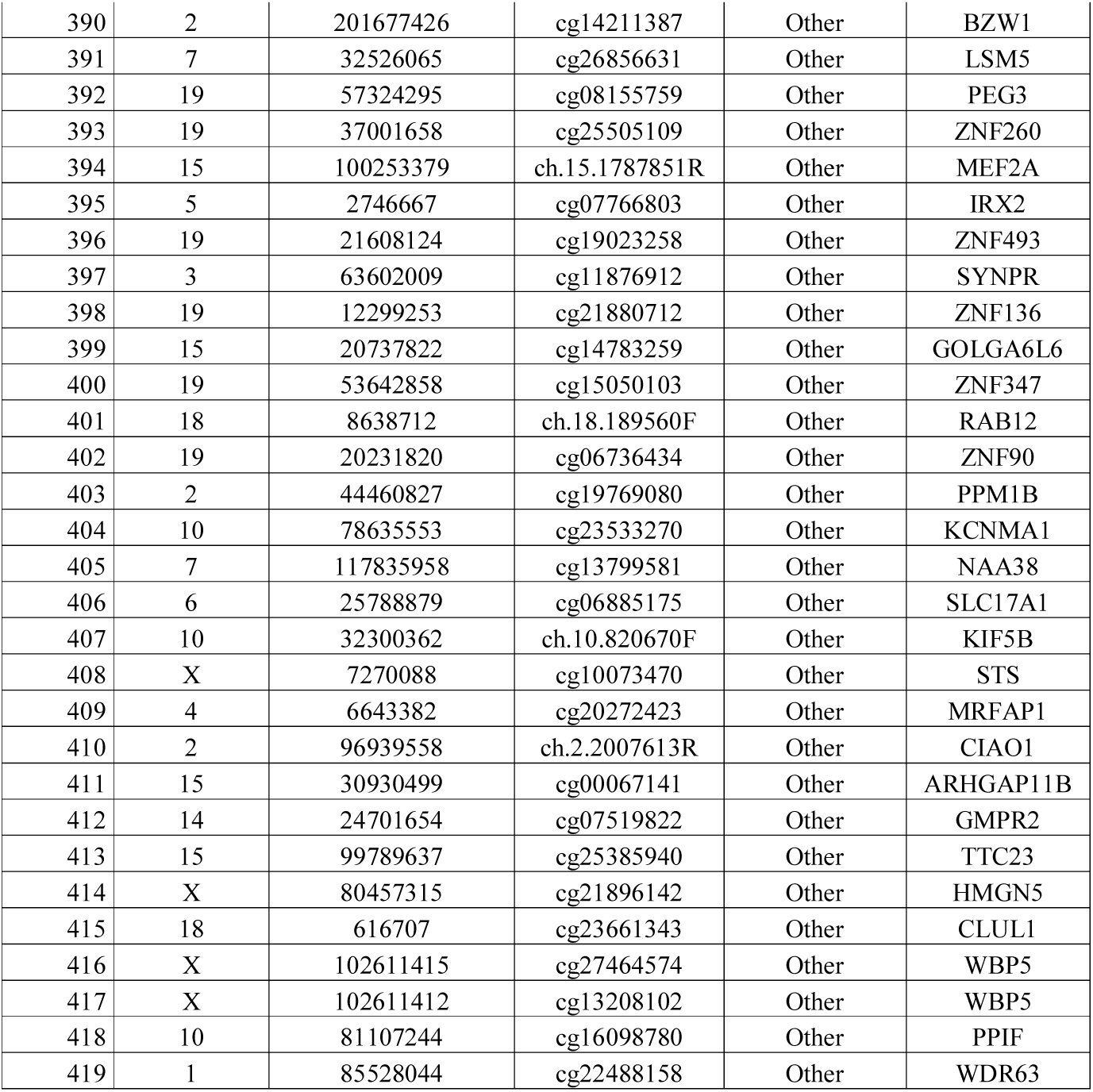
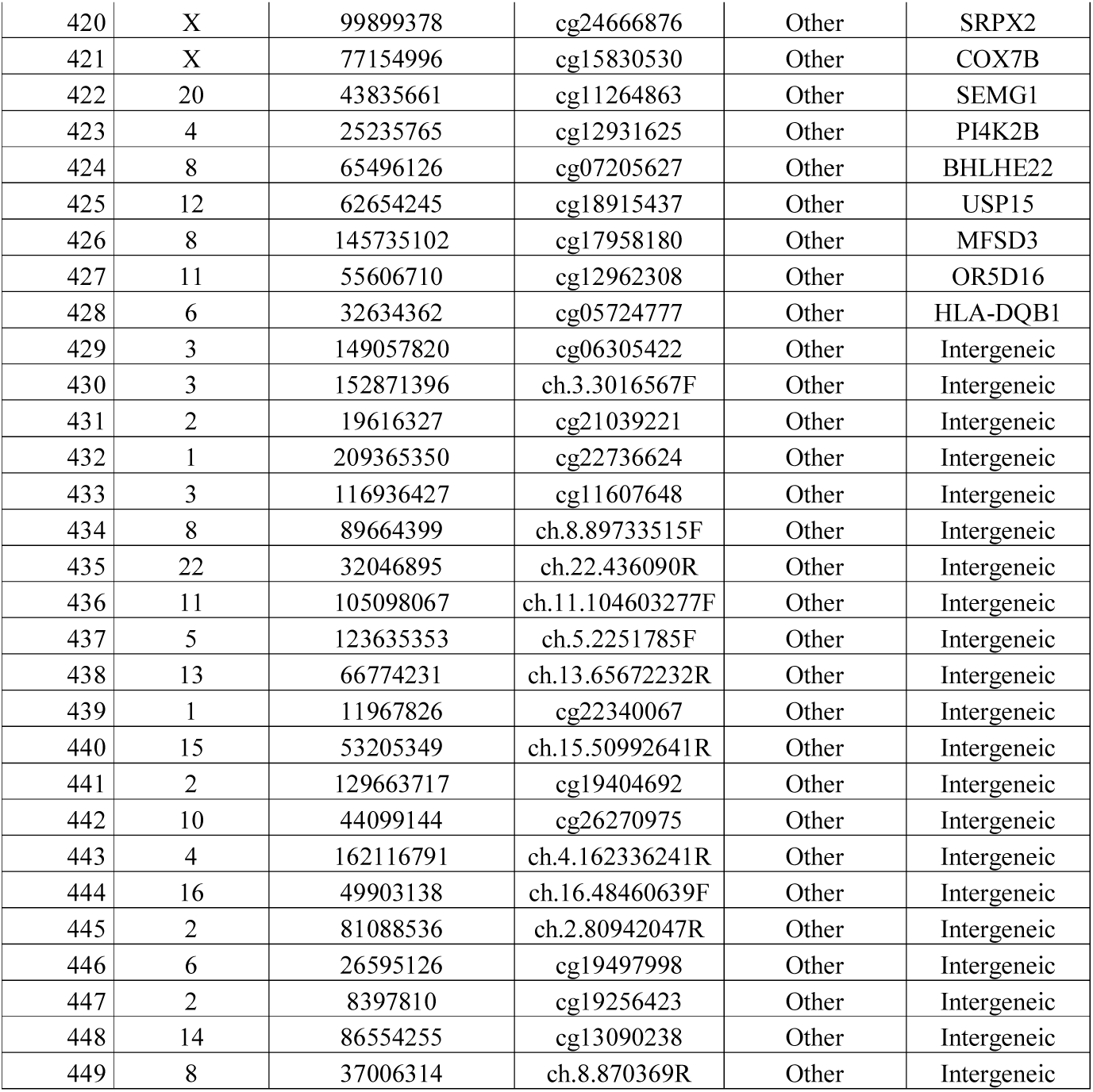
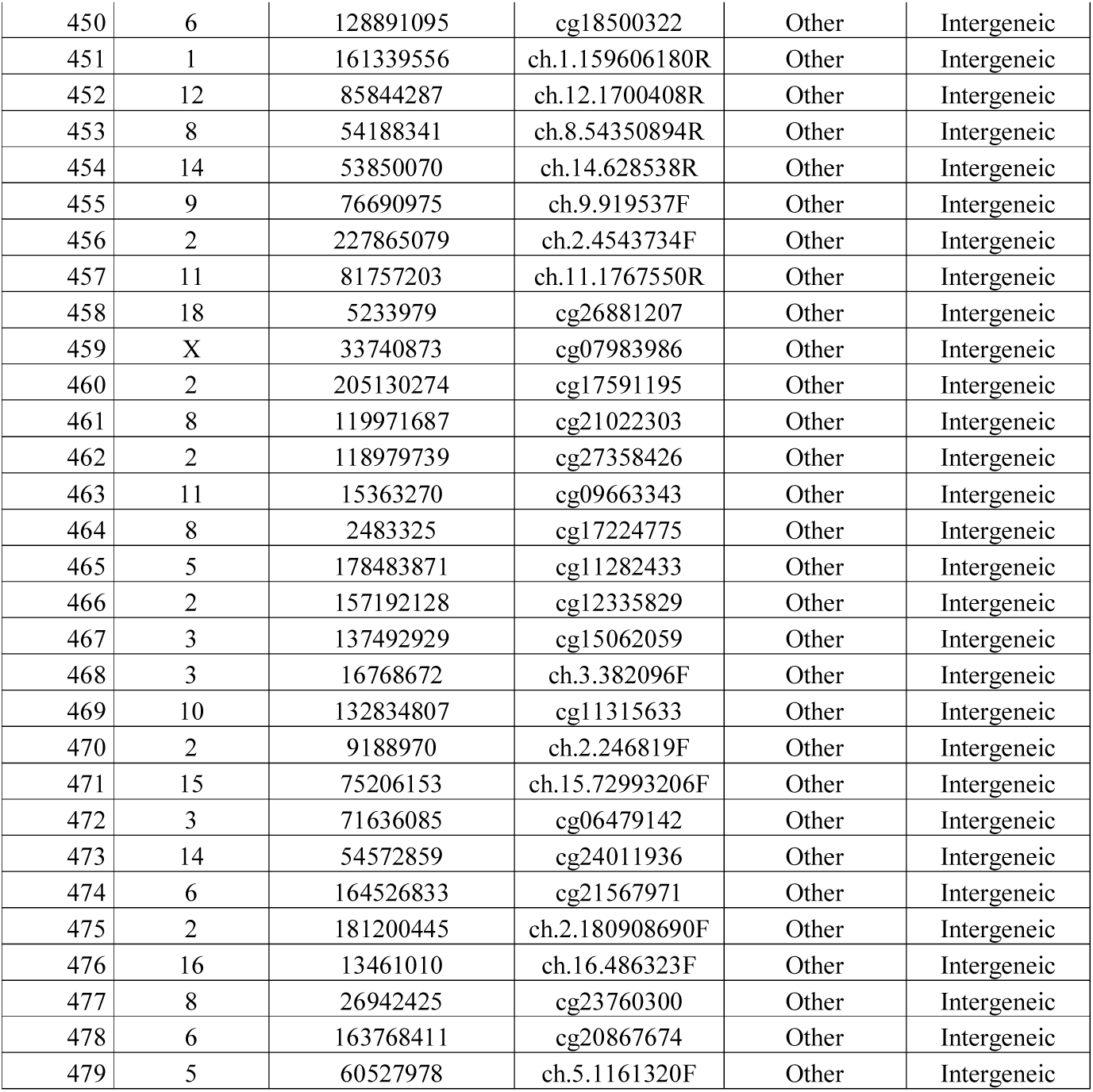
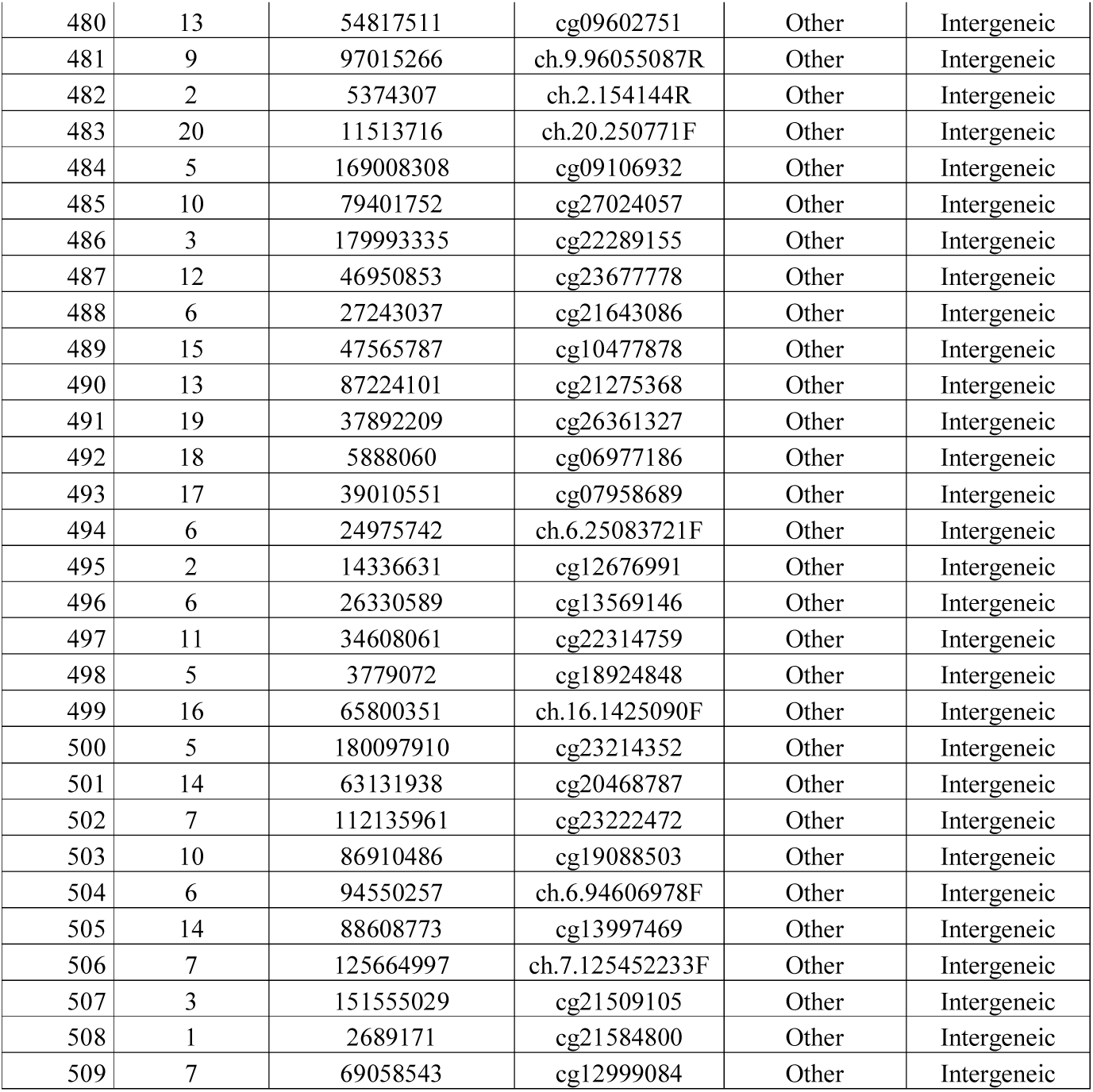
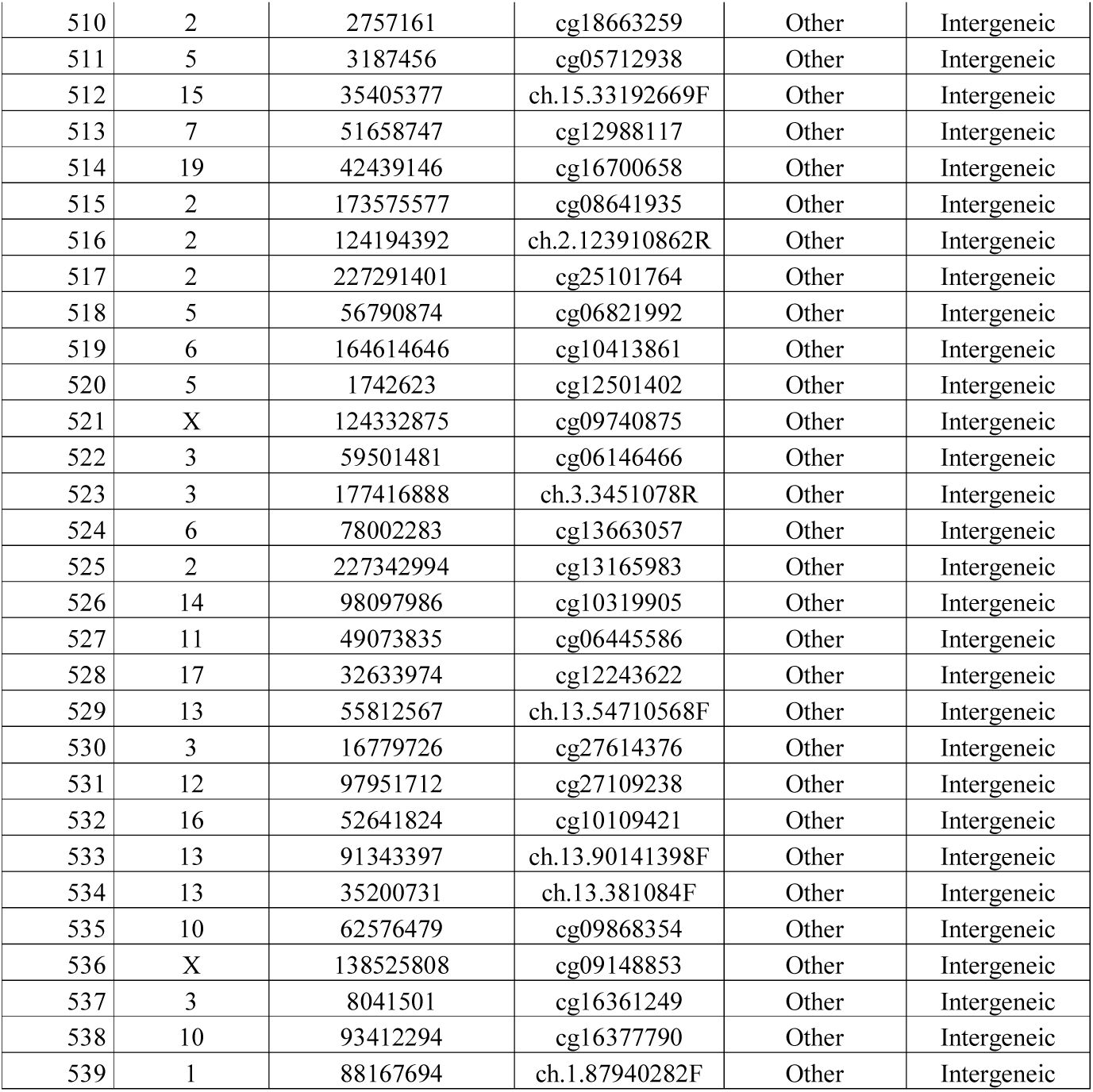
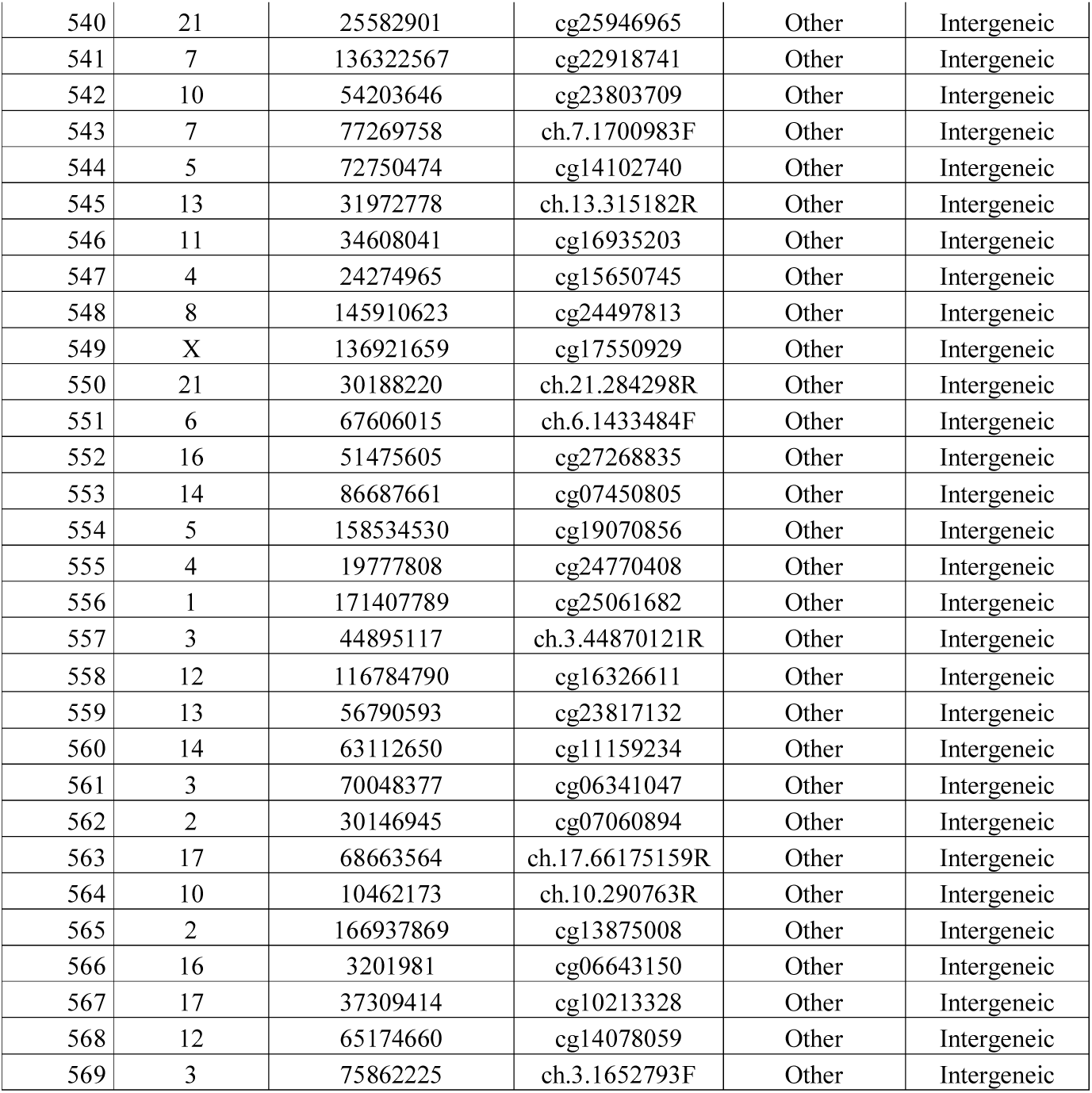
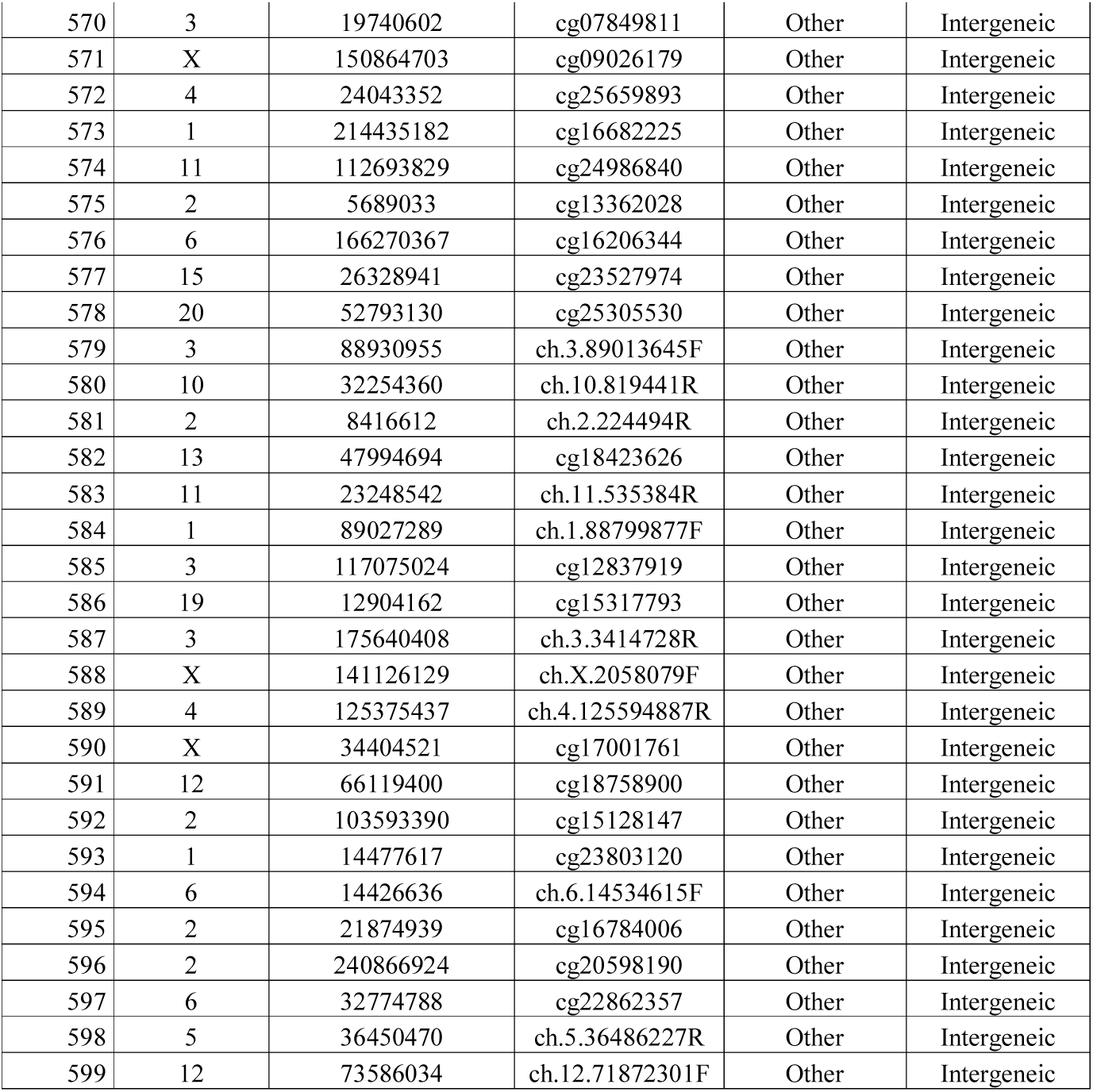
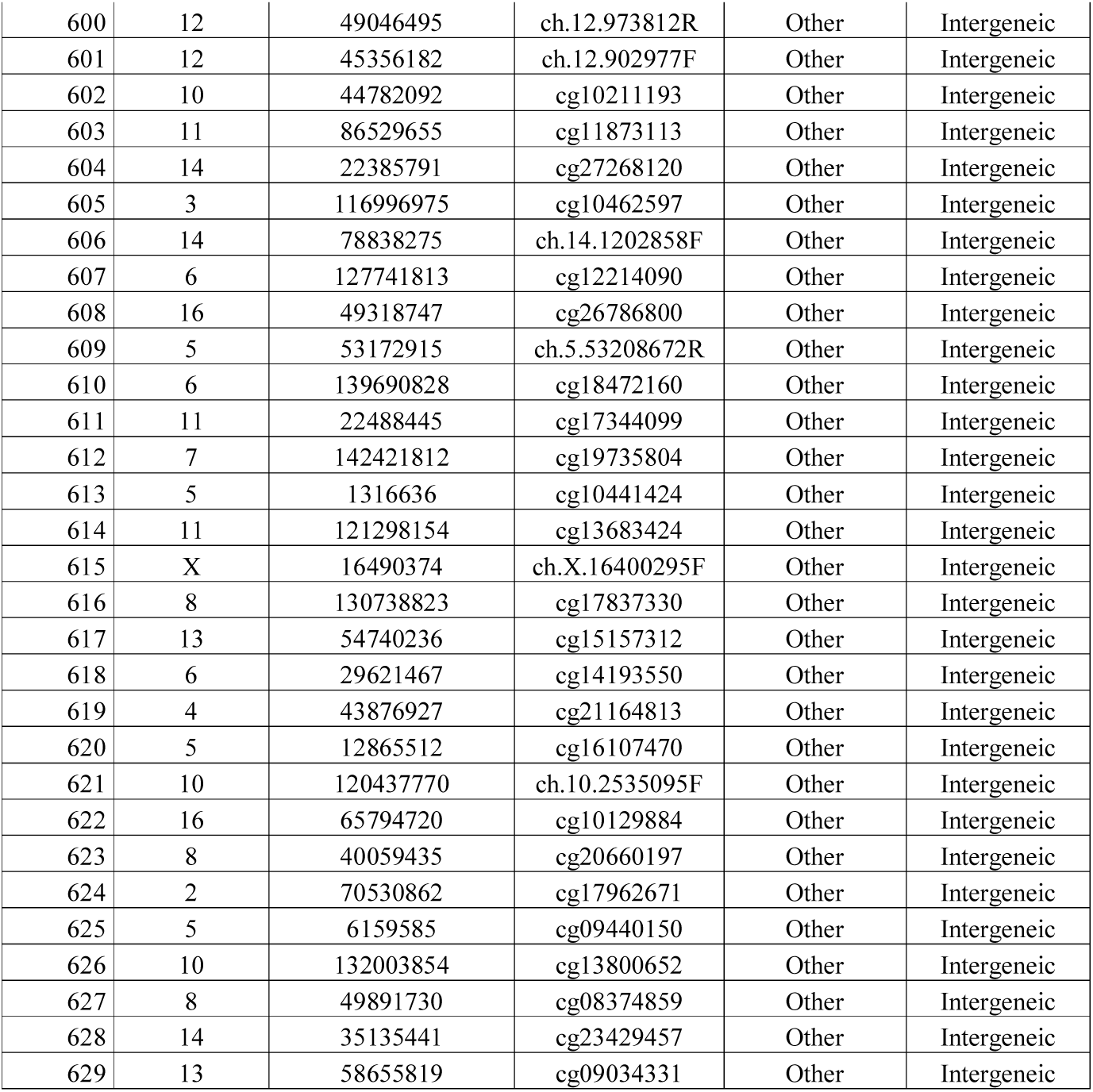
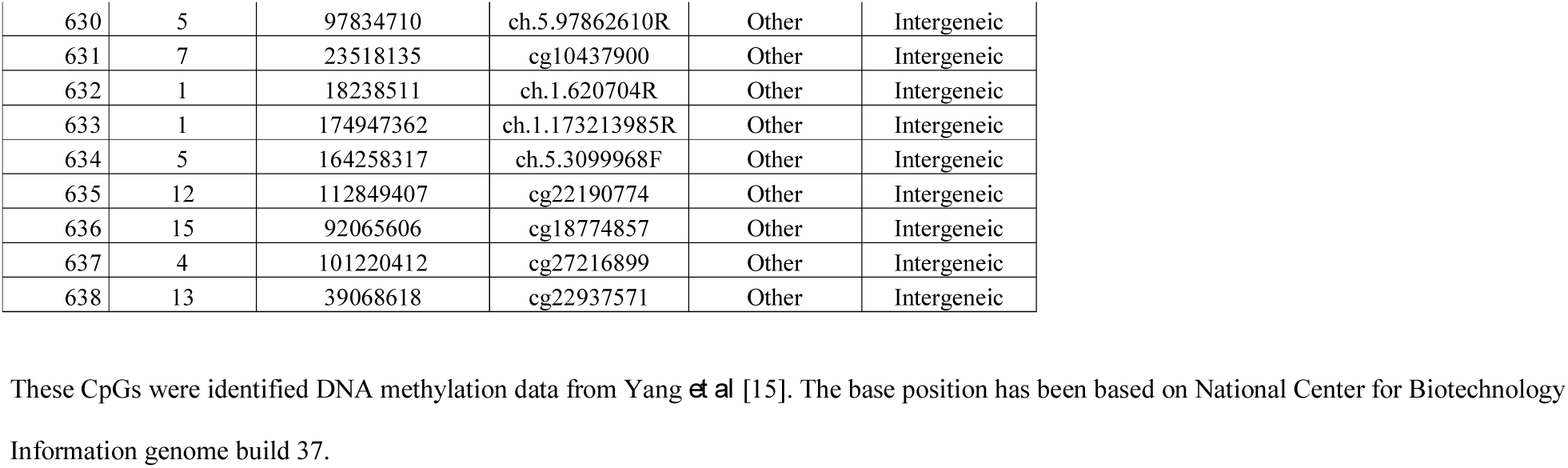
List of 638 CpG showing increased methylation after decitabine treatment in HCTK116 cell line

**Supplementary Table 2:**
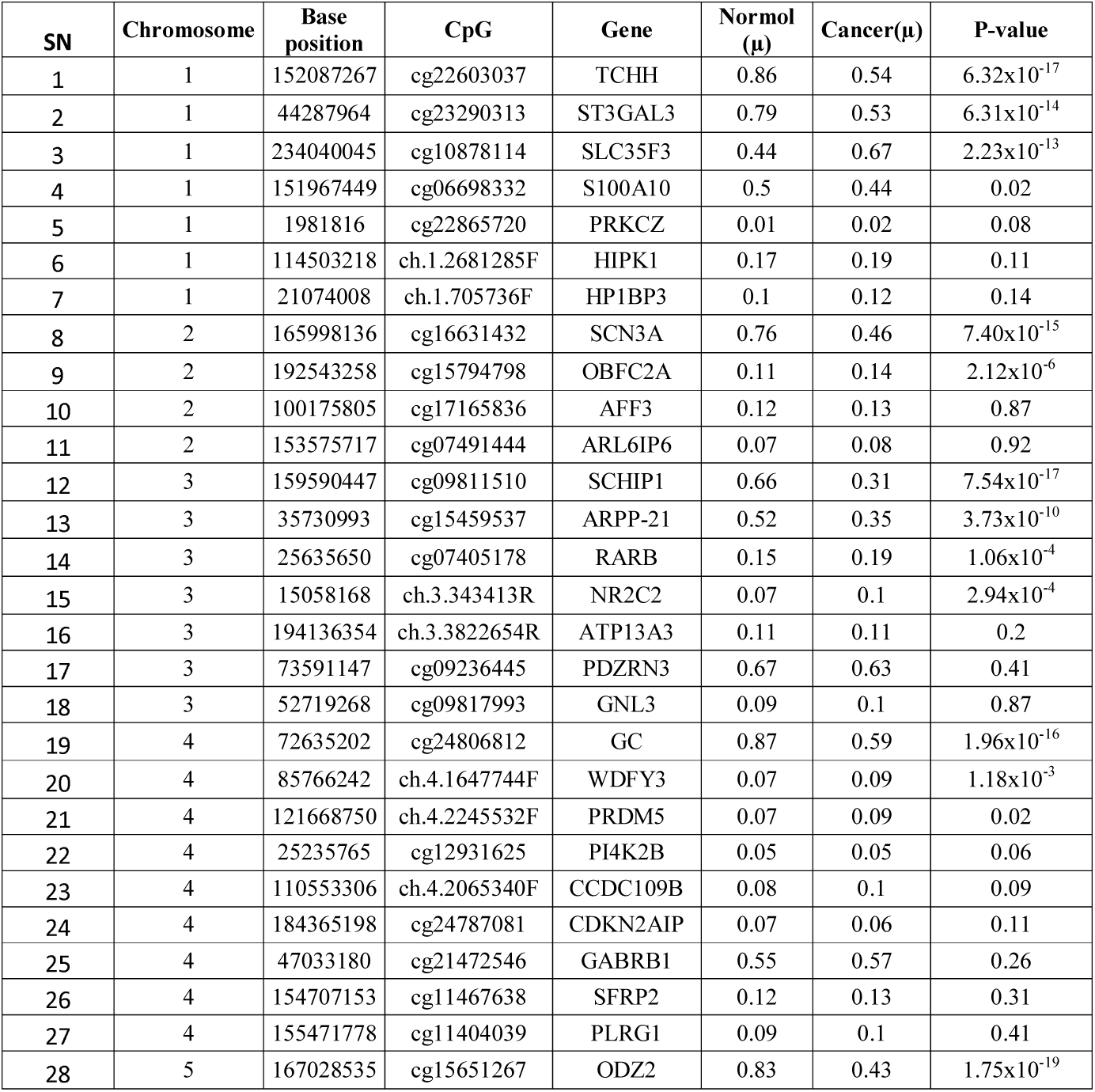
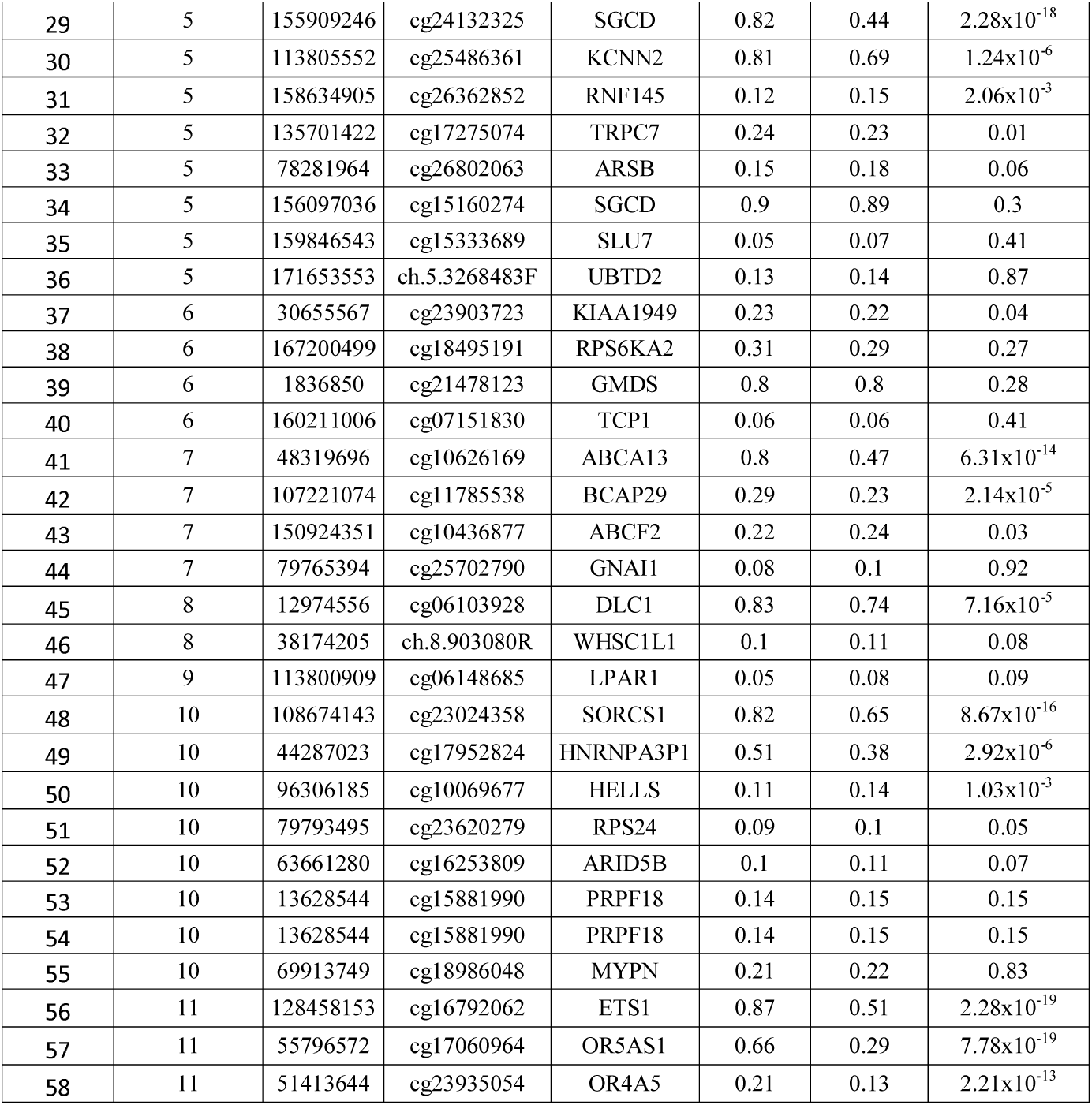
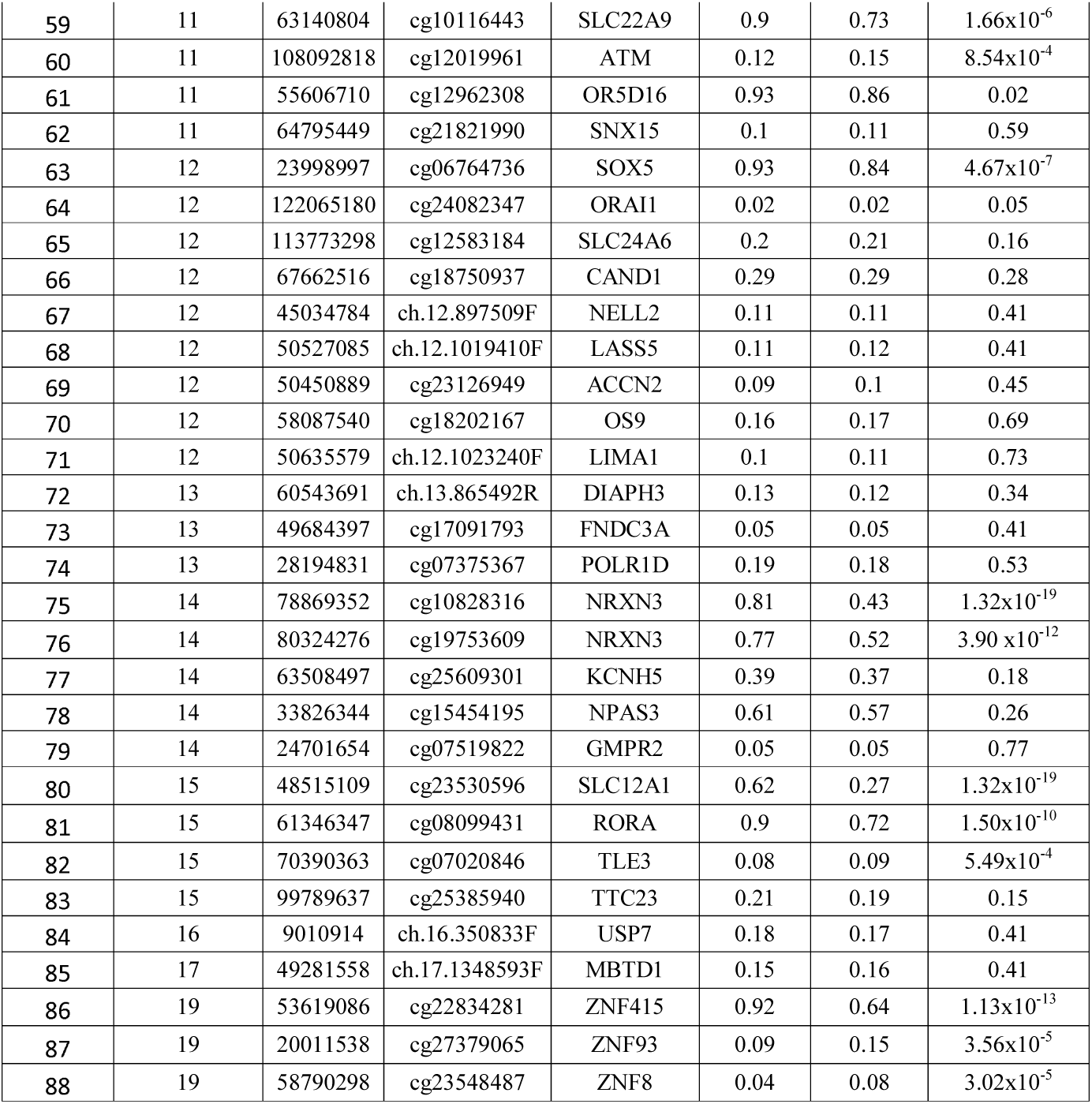
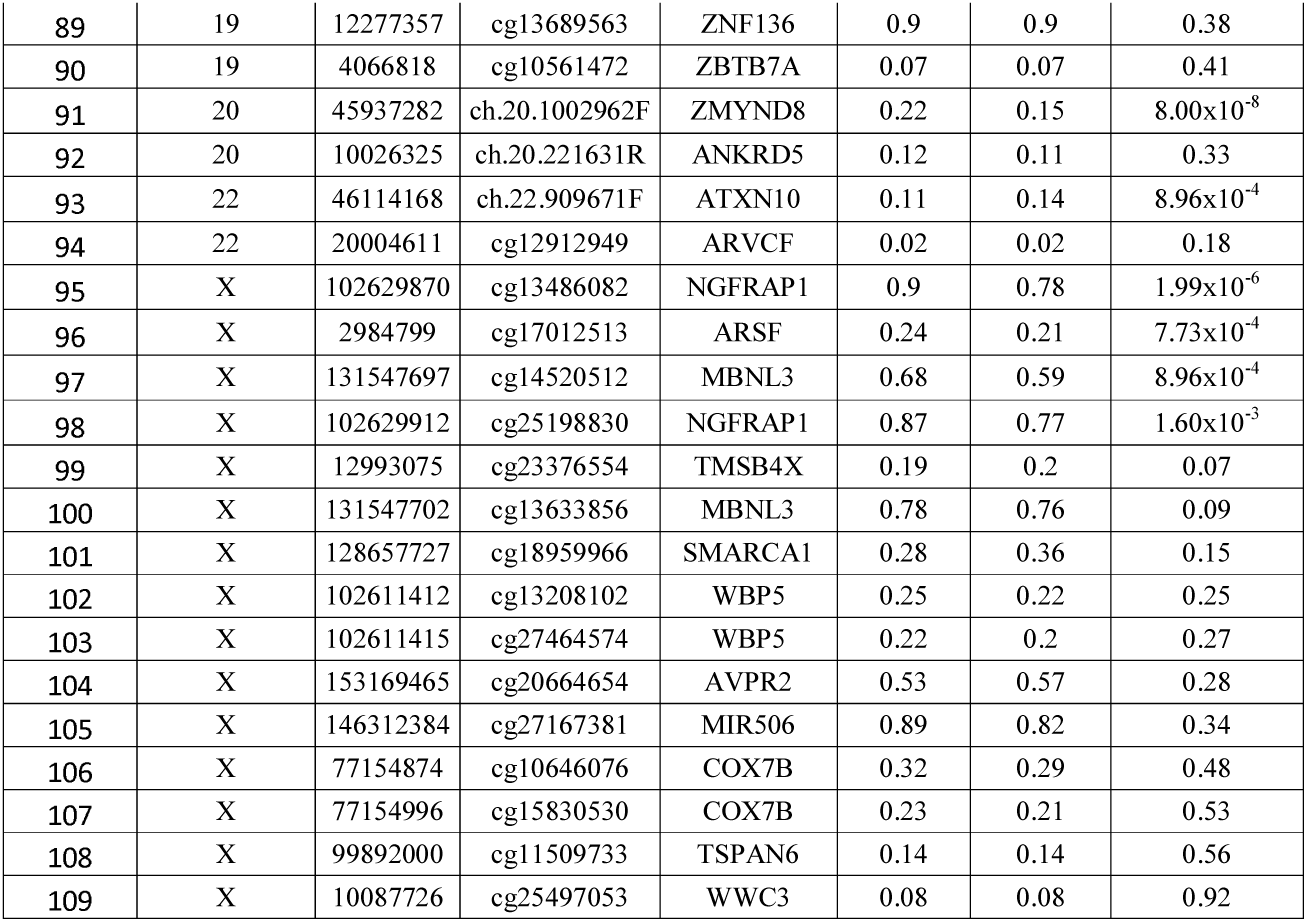
Differential methylation analysis of 109 identified CpGs in the TCGA colon cancer data

**Supplementary Table 3:** Differential expression analysis of identified probes in TCGA colon cancer data

| SN | Gene | log2(fold) | P-value |
| --- | --- | --- | --- |
| 1 | ABCF2 | -1.12 | 3.25x10<sup>-13</sup> |
| 2 | ACCN2 | -1.09 | 2.17x10<sup>-9</sup> |
| 3 | ARID5B | -1.28 | 1.35x10<sup>-3</sup> |
| 4 | ARSB | 1.06 | 0.27 |
| 5 | ARVCF | -1.05 | 1.22x10<sup>-8</sup> |
| 6 | ATM | 1.32 | 0.41 |
| 7 | AVPR2 | -1.05 | 3.00x10<sup>-7</sup> |
| 8 | BCAP29 | 1.03 | 1.76x10<sup>-3</sup> |
| 9 | CAND1 | -1.18 | 1.51x10<sup>-7</sup> |
| 10 | CDKN2AIP | -1.59 | 0.06 |
| 11 | COX7B | -1.25 | 9.78x10<sup>-3</sup> |
| 12 | ETS1 | 1.04 | 0.01 |
| 13 | FNDC3A | -1.15 | 0.06 |
| 14 | GABRB1 | -1.06 | 0.02 |
| 15 | GNL3 | -1.10 | 1.63x10<sup>-13</sup> |
| 16 | HNRNPA3P1 | -1.09 | 9.84x10<sup>-9</sup> |
| 17 | KIAA1949 | -1.41 | 0.33 |
| 18 | LPAR1 | -1.07 | 4.48x10<sup>-14</sup> |
| 19 | NGFRAP1 | -1.16 | 0.43 |
| 20 | NMT2 | -1.09 | 6.92x10<sup>-9</sup> |
| 21 | NRXN3 | -1.05 | 0.03 |
| 22 | NTM | -1.31 | 0.02 |
| 23 | OS9 | -1.13 | 5.76x10<sup>-3</sup> |
| 24 | PIGA | -1.12 | 5.85x10<sup>-7</sup> |
| 25 | PLRG1 | -1.35 | 0.01 |
| 26 | POLR1D | -1.08 | 3.25x10<sup>-13</sup> |
| 27 | PRKCZ | -1.30 | 0.06 |
| 28 | PRPF18 | -1.32 | 0.06 |
| 29 | RNF145 | -1.09 | 0.19 |
| 30 | RPS24 | -1.05 | $1.51 \times 10^{-7}$ |
| 31 | S100A10 | -1.07 | 0.01 |
| 32 | SEPSECS | -1.21 | 0.19 |
| 33 | SLC24A6 | -1.22 | $1.10 \times 10^{-6}$ |
| 34 | SLC35F3 | -1.05 | $1.81 \times 10^{-7}$ |
| 35 | SLU7 | -1.88 | 0.29 |
| 36 | SMARCA1 | -1.04 | $6.62 \times 10^{-3}$ |
| 37 | TCEAL8 | 1.00 | 0.76 |
| 38 | TCHH | -1.09 | 0.06 |
| 39 | TCP1 | -1.14 | $2.32 \times 10^{-7}$ |
| 40 | TLE3 | 1.06 | $7.74 \times 10^{-3}$ |
| 41 | TP73 | -1.14 | $4.28 \times 10^{-11}$ |
| 42 | TSPAN6 | -1.13 | $8.22 \times 10^{-3}$ |
| 43 | ZBTB7A | -1.13 | $4.25 \times 10^{-7}$ |
| 44 | ZNF8 | -1.05 | $3.36 \times 10^{-7}$ |
| 45 | ZNF93 | -1.02 | $2.40 \times 10^{-3}$ |
| 46 | ABCA13 | 1.13 | 0.59 |
| 47 | AFF3 | -1.09 | $3.25 \times 10^{-13}$ |
| 48 | AGTPBP1 | -1.07 | $1.99 \times 10^{-9}$ |
| 49 | ANKRD5 | -1.10 | $2.32 \times 10^{-7}$ |
| 50 | ARL6IP6 | -1.18 | $1.07 \times 10^{-9}$ |
| 51 | ATP13A3 | -1.12 | $2.17 \times 10^{-9}$ |
| 52 | ATXN10 | -1.19 | $4.74 \times 10^{-4}$ |
| 53 | CADM1 | -1.03 | $4.45 \times 10^{-6}$ |
| 54 | CCDC109B | -1.35 | $6.20 \times 10^{-5}$ |
| 55 | COX7B1 | -1.25 | $9.78 \times 10^{-3}$ |
| 56 | CUL3 | -1.07 | 0.46 |
| 57 | DIAPH3 | -1.11 | $1.76 \times 10^{-13}$ |
| 58 | DLC1 | -1.01 | $7.01 \times 10^{-4}$ |
| 59 | FAT3 | -1.03 | 0.06 |
| 60 | GAB2 | -1.11 | $3.08 \times 10^{-3}$ |
| 61 | GMDS | -1.04 | 0.02 |
| 62 | GMPR2 | -1.02 | 0.08 |
| 63 | GNAI1 | -1.08 | $1.32 \times 10^{-9}$ |
| 64 | HELLS | -1.11 | $2.49 \times 10^{-13}$ |
| 65 | HIPK1 | 1.00 | 0.10 |
| 66 | HP1BP3 | -1.19 | 0.82 |
| 67 | KCNN2 | 1.27 | 0.34 |
| 68 | LARS2 | -1.12 | $6.61 \times 10^{-5}$ |
| 69 | LASS5 | -1.09 | $8.59 \times 10^{-9}$ |
| 70 | LIMA1 | -1.10 | $1.11 \times 10^{-10}$ |
| 71 | MAP7 | -1.11 | $6.43 \times 10^{-7}$ |
| 72 | MBNL3 | -1.06 | $2.26 \times 10^{-5}$ |
| 73 | MBTD1 | -1.12 | $1.76 \times 10^{-3}$ |
| 74 | MRPS9 | 1.05 | 0.04 |
| 75 | MYPN | -1.12 | $8.16 \times 10^{-10}$ |
| 76 | NELL2 | -1.11 | $4.36 \times 10^{-7}$ |
| 77 | NPAS3 | -1.08 | $1.90 \times 10^{-10}$ |
| 78 | NR2C2 | -1.27 | $9.78 \times 10^{-3}$ |
| 79 | NRXN31 | -1.05 | 0.03 |
| 80 | OBFC2A | -1.03 | $3.08 \times 10^{-3}$ |
| 81 | ODZ2 | -1.04 | $1.50 \times 10^{-6}$ |
| 82 | ORAI1 | -1.09 | $1.15 \times 10^{-4}$ |
| 83 | PDE1C | -1.10 | $6.74 \times 10^{-12}$ |
| 84 | PDZRN3 | -1.09 | $5.34 \times 10^{-8}$ |
| 85 | PI4K2B | -1.16 | $5.32 \times 10^{-7}$ |
| 86 | PRDM5 | -1.12 | 0.90 |
| 87 | RARB | 1.11 | 0.24 |
| 88 | RFC3 | -1.10 | $8.98 \times 10^{-14}$ |
| 89 | RGMB | -1.13 | $8.22 \times 10^{-3}$ |
| 90 | RORA | -1.03 | $7.16 \times 10^{-5}$ |
| 91 | RPS6KA2 | -1.17 | 0.24 |
| 92 | SCHIP1 | -1.04 | $1.70 \times 10^{-4}$ |
| 93 | SCN3A | -1.08 | $1.76 \times 10^{-13}$ |
| 94 | SFRP2 | 1.00 | $3.39 \times 10^{-5}$ |
| 95 | SGCD | 1.01 | 0.01 |
| 96 | SMYD4 | -1.08 | $5.76 \times 10^{-3}$ |
| 97 | SNX15 | -1.10 | $1.42 \times 10^{-4}$ |
| 98 | SORCS1 | -1.08 | $3.39 \times 10^{-13}$ |
| 99 | SOX5 | -1.10 | $2.32 \times 10^{-7}$ |
| 100 | ST3GAL3 | -1.09 | $1.04 \times 10^{-7}$ |
| 101 | TRNAU1AP | 1.14 | 0.52 |
| 102 | TTC23 | -1.15 | $2.74 \times 10^{-6}$ |
| 103 | UBE2L3 | -22.63 | 0.45 |
| 104 | UBTD2 | -1.06 | $2.49 \times 10^{-3}$ |
| 105 | USP7 | -1.07 | $2.32 \times 10^{-4}$ |
| 106 | WBP5 | -1.21 | 0.45 |
| 107 | WDFY3 | 1.14 | 0.52 |
| 108 | WHSC1L1 | -1.22 | 0.07 |
| 109 | WWC3 | -1.09 | $3.52 \times 10^{-5}$ |
| 110 | ZMYND8 | -1.09 | $2.90 \times 10^{-6}$ |
| 111 | ZNF136 | 3.17 | 0.59 |
| 112 | ZNF415 | -1.09 | $5.58 \times 10^{-9}$ |

**Supplementary figure 1:**
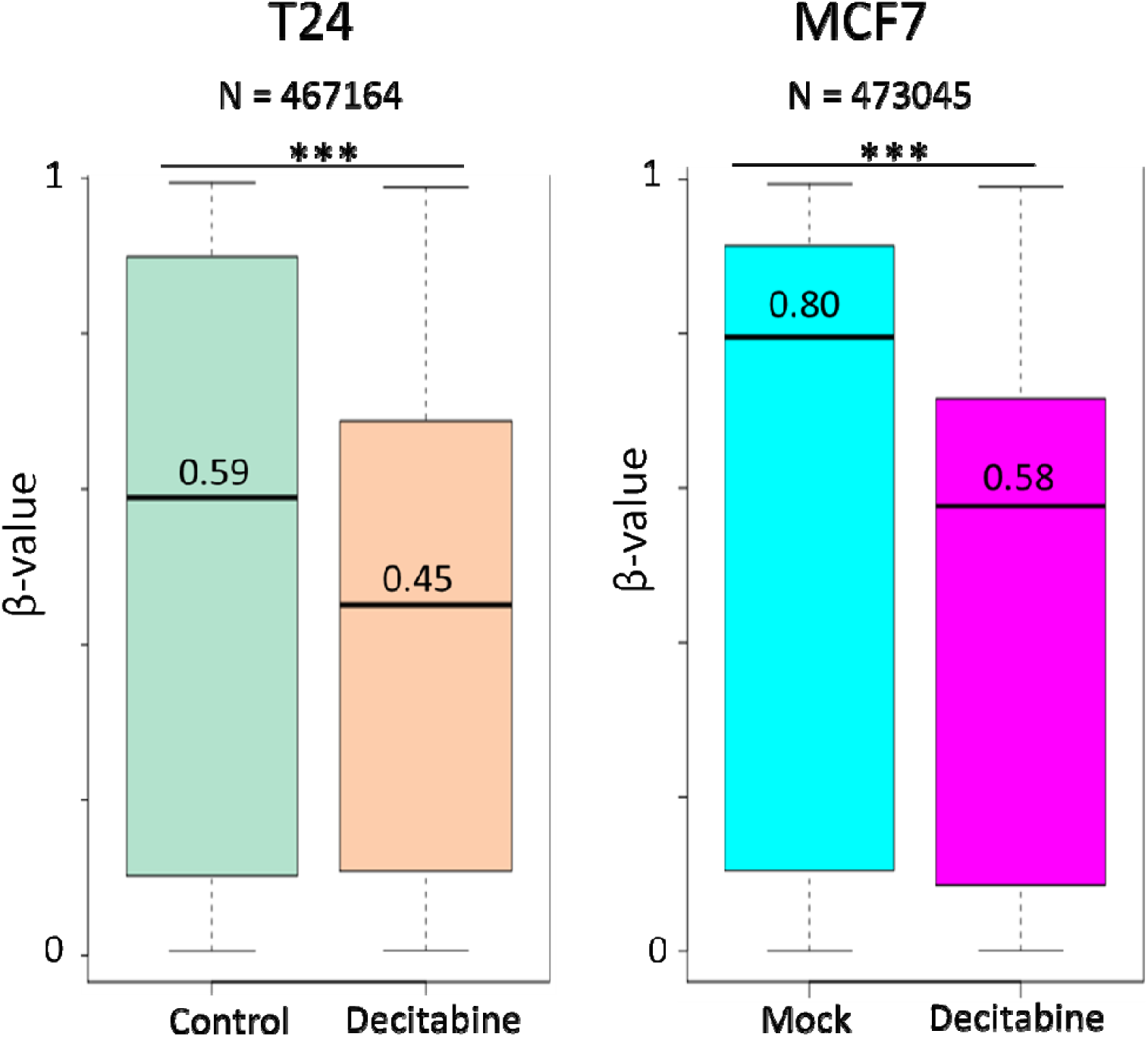
Box plot showing a decrease in methylation level of loci other than identified CpGs in T24 and MCF7 cell line after decitabine treatment. N denotes the total number of CpGs analyzed in the data. The data from the study by Han *et al* [16] (GSE41525) and Leadem *et al* [17] (GSE97483) has been shown for T24 and MCF7 cells respectively

**Supplementary figure 2:**
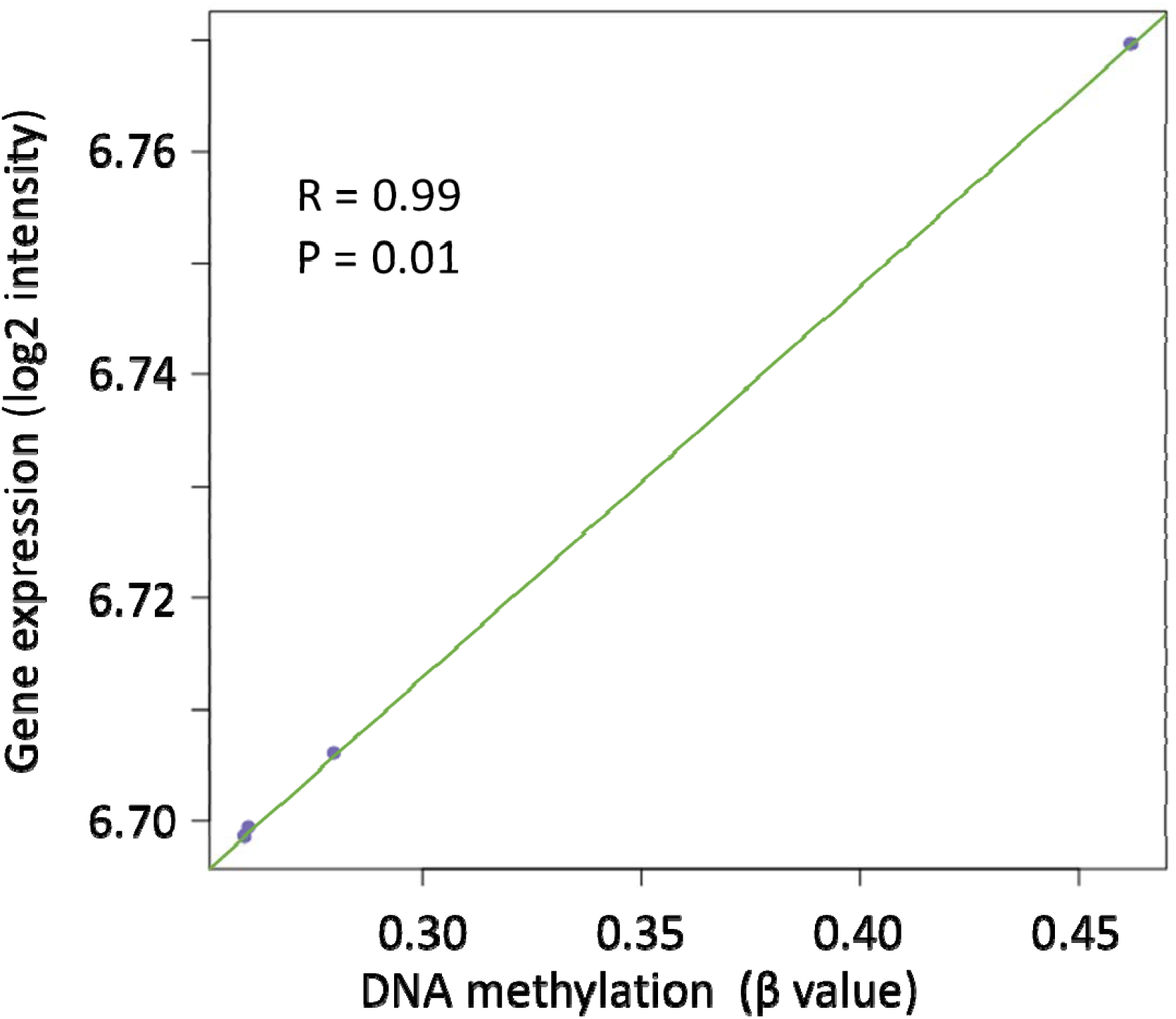
Scatter plot showing the correlation between RORA expression level and methylation level at CpGs cg08099431 in HCT116 cell line.

**Supplementary figure 3:**
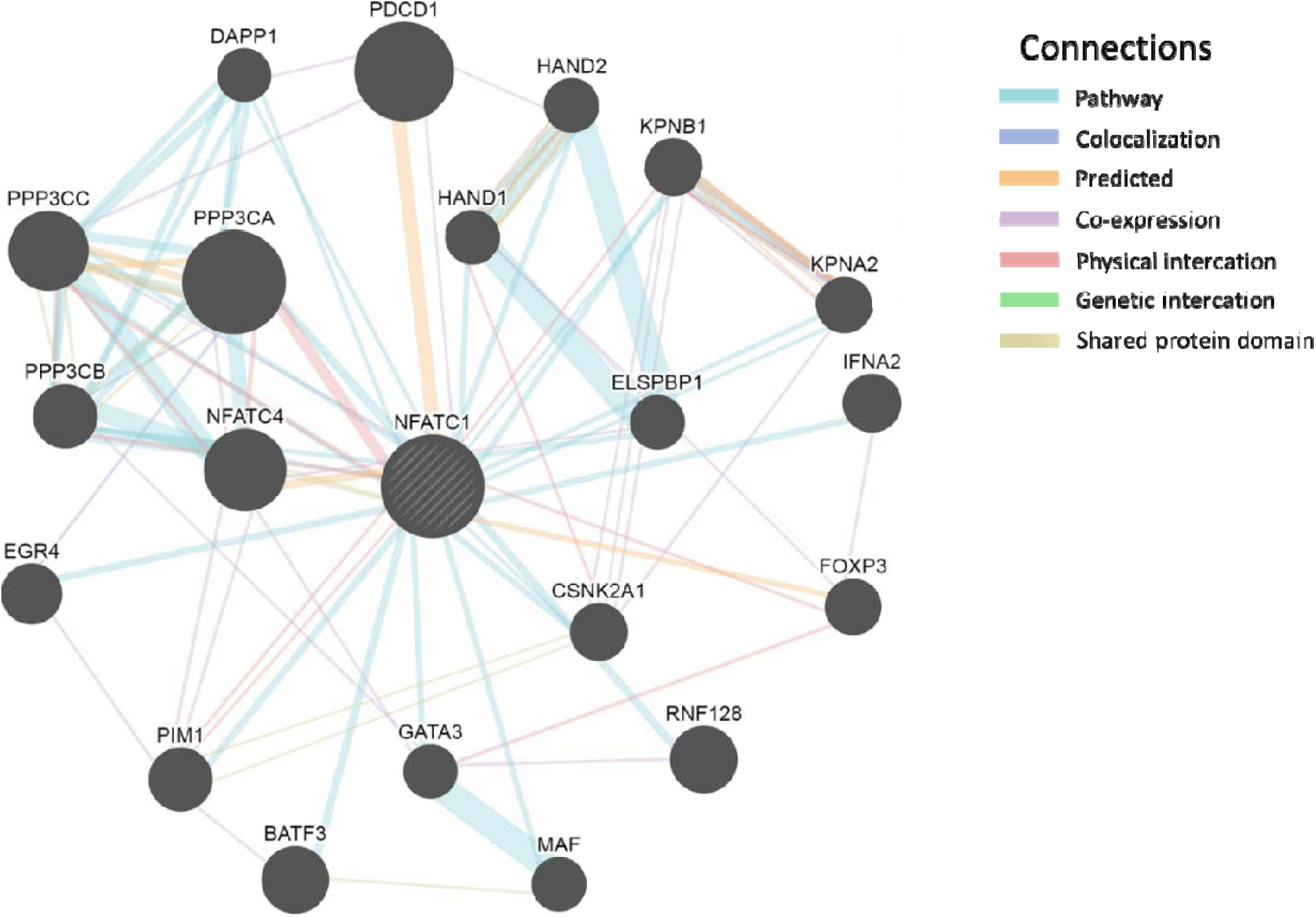
Interaction network of genes regulated by NFATc1. Only interaction among those genes that are directly connected to NFATc1 has been shown using network construction from GENEMANIA. The size of the gene nodes is proportional to gene score calculated by GENEMANIA using label propagation algorithm that indicates the relevance of each gene to the original list based on the selected networks.

**Supplementary figure 4:**
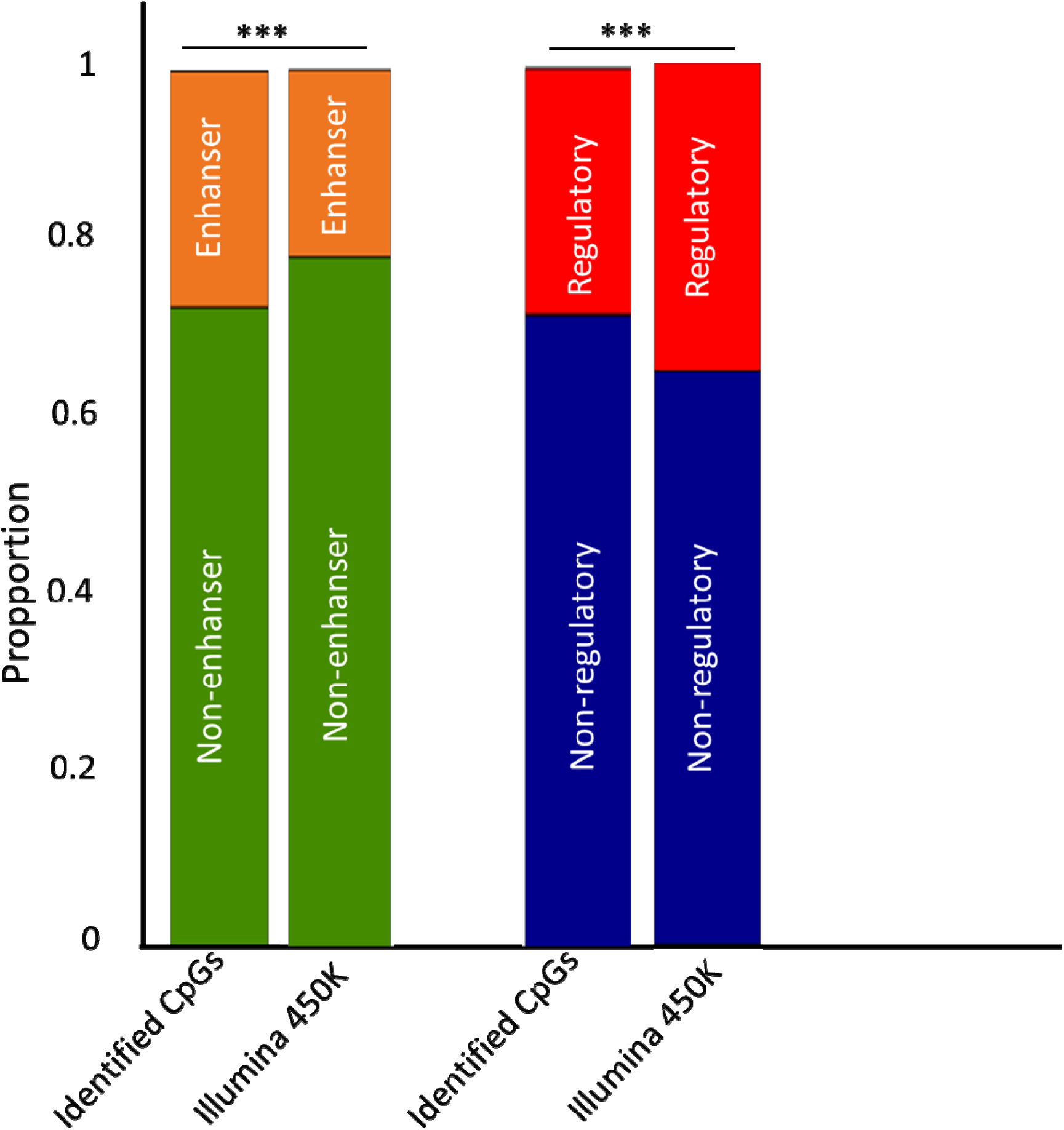
Enrichment analysis of identified CpGs in enhancer and regulatory region Eenrichment of identified CpGs among regulatory region of the genome. CpGs with increased methylations were enriched in enhancer region (21% of identified CpGs were in enhancer region as compared to 26% of total CpGs present in 450K chip, P = 3.36x10^−4^) and were depleted in other regulatory regions such as promoter and non-promoter associated cell type-specific or general regulatory regions represented by transcription factor binding sites and DNA hypersensitivity elements (25% of identified CpGs were in other regulatory region as compared to 28% of total CpGs in 450K beadchip, P = 2.65x10^−4^). The proportions of identified CpGs in enhancer and regulatory region have been shown as segmented barplot (upper segment). ***P<0.0005

**Author contribution:** AKG conceptualized, analyzed the data and wrote the manuscript. TA critically revised and edited the manuscript. The authors report no conflict of interest

